# Continuous sumatriptan exposure induces persistent trigeminovascular sensitisation and brain perfusion changes in a rat model of medication overuse headache

**DOI:** 10.64898/2026.05.26.726530

**Authors:** James GA Hall, Fiona M Boissonade, Aneurin J Kennerley, Milena De Felice

## Abstract

**Background:** Repeated exposure to acute antimigraine medication can promote medication overuse headache, but the mechanisms underlying this transition remain incompletely understood. We used a clinically relevant rat model of continuous sumatriptan exposure to investigate whether medication overuse is associated with persistent sensitisation of the trigeminovascular system and longer-lasting changes in brain perfusion.

**Methods:** Adult male Sprague Dawley rats received continuous subcutaneous sumatriptan (0.6 mg/kg/day) or saline infusion for 6 days via osmotic minipumps. Periorbital and hindpaw mechanical thresholds were measured over 20 days. On day 6 and day 20, trigeminal ganglia and trigeminal nucleus caudalis were processed for immunohistochemistry for pERK, pp38, Iba-1, GFAP and NeuN. On day 20, a subgroup received sodium nitroprusside (SNP, 3 mg/kg, i.p.) to unmask latent sensitisation. Cerebral blood flow was assessed by MRI.

**Results:** Sumatriptan induced reversible cephalic and extracephalic allodynia. Previously exposed rats showed evidence of persistent sensitisation, including enhanced biomarker and glial responses after withdrawal and following SNP challenge. pERK and pp38 expression increased in both the trigeminal ganglion and trigeminal nucleus caudalis. In the TNC, marker association shifted over time from predominantly neuronal at day 6 to greater apparent glial association at day 20. Iba-1 and GFAP expression increased after withdrawal of sumatriptan and was further enhanced by SNP challenge. Within the TNC, neuronal marker expression was greatest in the ophthalmic representation. Sumatriptan exposure also produced a persistent reduction in cerebral blood flow that remained evident after behavioural recovery.

**Conclusion:** Continuous sumatriptan exposure produces prolonged trigeminovascular neuronal and glial alterations together with persistent changes in brain perfusion. These data support a state of latent sensitisation after repeated triptan exposure and provide mechanistic insight into medication overuse headache.

**HIGHLIGHTS:**

- Repeated sumatriptan exposure induces reversible cephalic and extracephalic allodynia but leaves persistent trigeminovascular sensitisation after drug withdrawal.
- pERK and pp38 expression increase in the trigeminal ganglion and trigeminal nucleus caudalis, with the strongest regional changes seen in the ophthalmic representation of the TNC.
- Delayed increases in Iba-1 and GFAP in the TNC suggest that glial activation may contribute to maintenance of latent sensitisation, although the colocalisation findings are qualitative and should be interpreted cautiously.
- Repeated sumatriptan exposure is also associated with a persistent reduction in cerebral blood flow, indicating longer-lasting changes in brain perfusion beyond the period of overt allodynia.

## INTRODUCTION

Migraine is an incapacitating collection of neurological symptoms that usually results in a severe, throbbing, recurring pain on one side of the head. Migraine is the most common neurological condition in developed countries. The World Health Organisation ranks migraine the third most common disease globally and the second leading cause of years lived with disability (1, 2).

According to the Global Burden of Disease Study in 2019, migraine affects 1.1 billion people worldwide, including about 8 million people in the UK alone. Its incidence is rising in many parts of the world. In combination with the absence of effective long-term and/or acute treatments, it presents a significant global health concern. Migraine is a source of significant pain and can cause incapacitation for days after an attack. Although it is a common disease, migraine’s pathophysiology is poorly understood. Trigeminovascular-mediated central sensitisation has been implicated in the development of migraine symptoms, including pain following light touch (cutaneous allodynia). Many but not all patients, report cutaneous allodynia during migraine, with a distribution that can include both cephalic and extracephalic areas (3, 4). Central sensitisation, the presumed underlying physiological substrate of cutaneous allodynia, is thought to underlie the progression of migraine pain, and possibly the transformation from an episodic to a chronic, more disabling, disorder known as chronic migraine (5–7). A major role in the shift from episodic to chronic migraine in a significant proportion of patients is dictated by the overuse of medications, including triptans, that are originally prescribed for migraine itself (6, 8–13), and can lead to a syndrome called medication overuse headache (MOH) (14–17).

Triptans are thought to relieve headache pain by inhibiting the release of pronociceptive transmitters via actions on 5HT1B/1D receptors found on trigeminal afferent neurones (18). However, whilst they elicit important therapeutic actions, triptans can also produce a maladaptive response that increases the occurrence of migraine and can promote chronic migraine and MOH (19–21). Multiple preclinical observations indicate that repeated exposure to triptans induces persistent neural adaptations within trigeminovascular pathways, producing a state of latent sensitisation that enhances responsiveness to known migraine triggers and may contribute to the development of chronic migraine and medication overuse headache (22–24). However, the precise mechanisms driving MOH in humans remain incompletely understood. Thus, investigation of the mechanisms that contribute to central sensitisation is of considerable relevance to our understanding of migraine pain and chronification of migraine, and will aid the development of novel therapeutic targets.

Migraine is increasingly recognised as a disorder of neuronal dysfunction involving maladaptive interactions between neurons and non-neuronal cells within pain-processing pathways (25, 26). While human studies have identified alterations in multiple brain regions associated with migraine, including those involved in nociceptive modulation and autonomic regulation ((i.e. hippocampus, basal ganglia (27, 28); medial brainstem and cingulate, auditory and visual cortices (29); periaqueductal gray, nucleus cuneiformis (30,31,32); autonomic function (33)). Such observations provide limited mechanistic insight into how repeated exposure to migraine medications drives sustained sensitisation. Preclinical approaches therefore remain essential for dissecting the cellular substrates of migraine-relevant plasticity within defined trigeminovascular circuits.

One approach to furthering our understanding of migraine pathology is to use specific animal models. We have previously reported that treatment with triptans over a period of days elicits maladaptations that offer opportunities to explore the biology of migraine and of MOH/chronic migraine (23, 24, 34, 35). In this model continuous infusion of sumatriptan for 7 days results in a state of cutaneous allodynia that slowly resolves within 14 days after stopping sumatriptan administration. A period of “latent sensitisation” follows (i.e. day 20), which is characterized by increased sensitivity to putative migraine triggers in humans including nitric oxide (NO) donors or environmental stress; increased expression of calcitonin gene-related peptide (CGRP) and nNOS, but not substance P, in identified dural afferents; and increased levels of CGRP in the jugular blood. Importantly it has been shown that mechanisms known to relieve migraine in humans reverse both cephalic and extracephalic allodynia (7,23,24,35). Furthermore, other studies have indicated a key role for the peripheral C-C motif ligand 2 (CCL2)-C-C motif chemokine receptor 2 (CCR2) pathways. The CCL2-it aids migraine resolution through low-dose interleukin-2 (LD-IL-2) by enhancing Treg cell function (36,37). Here we report that exposure of uninjured rats to triptans over several days leads to the development of both cephalic and extra-cephalic cutaneous allodynia, accompanied by enhanced neuronal activity within the trigeminovascular system, that persists following discontinuation of triptan administration. This sustained neuronal activation within the trigeminal nucleus caudalis, and trigeminal ganglion is likely to contribute to the behavioural and neurochemical hypersensitivity observed in response to stimuli known to trigger migraine in humans, despite otherwise normal baseline sensory thresholds. Such persistent sensitisation of trigeminovascular pathways may provide a mechanistic basis for medication-overuse headache, resulting in a lowered threshold for migraine initiation following repeated triptan exposure.

## MATERIALS AND METHODS

### Animals

Twenty-four adult male Sprague Dawley rats (175-250g) were maintained in a climate-controlled room on a 12 hr light/dark cycle with food and water ad libitum. All animal procedures were authorized by the UK Home Office and were performed in accordance with the UK Home Office Animals (Scientific Procedures) Act, 1986, and the policies and recommendations of the International Association for study of Pain and the National Institutes of Health guidelines for the handling and use of laboratory animals. Groups of 4 animals were used in all experiments. The ARRIVE guidelines have been followed in reporting this study.

### Surgery and drug administration

Continuous drug infusion was performed using Alzet osmotic minipumps (model 2001; 1 µl/hr/7 days; subcutaneous). On day 0, after evaluation of periorbital and hindpaw sensory threshold, rats underwent anaesthesia (isoflurane; 4% induction, 1.5% maintenance) and were subcutaneously implanted with one osmotic minipump containing either sumatriptan (AbMole, 0.6 mg/kg/day, dissolved in saline) or saline (1ml/kg/day). Rats were allowed to recover for 2 days, before periodic testing for allodynia was performed. At day 20 post implant of osmotic minipump and after evaluation of sensory threshold, some rats (as detailed below) were administered with sodium nitroprusside (SNP, 3 mg/kg, i.p., dissolved in saline), to induce a migraine-like state in the animals previously exposed to sumatriptan.

Rats were randomly assigned to receive sumatriptan (N=12) or saline (N=12). On day 6, 4 of the saline and 4 of the sumatriptan rats were deeply anaesthetised with an overdose of pentobarbitone for tissue collection. The remaining 16 rats had their minipumps removed and continued to be periodically tested for allodynia. On day 20, after testing of allodynia, half of the rats (4 from the sumatriptan and 4 from the saline pre-exposed group) were deeply anaesthetised with an overdose of pentobarbitone for tissue collection. The other half (4 from the sumatriptan and 4 from the saline pre-exposed group) received sodium nitroprusside (SNP, 3 mg/kg, i.p.), were tested for allodynia at 1h post-SNP injection, and then were deeply anaesthetised with an overdose of pentobarbitone for tissue collection (Figure 1).

**Figure 1.**
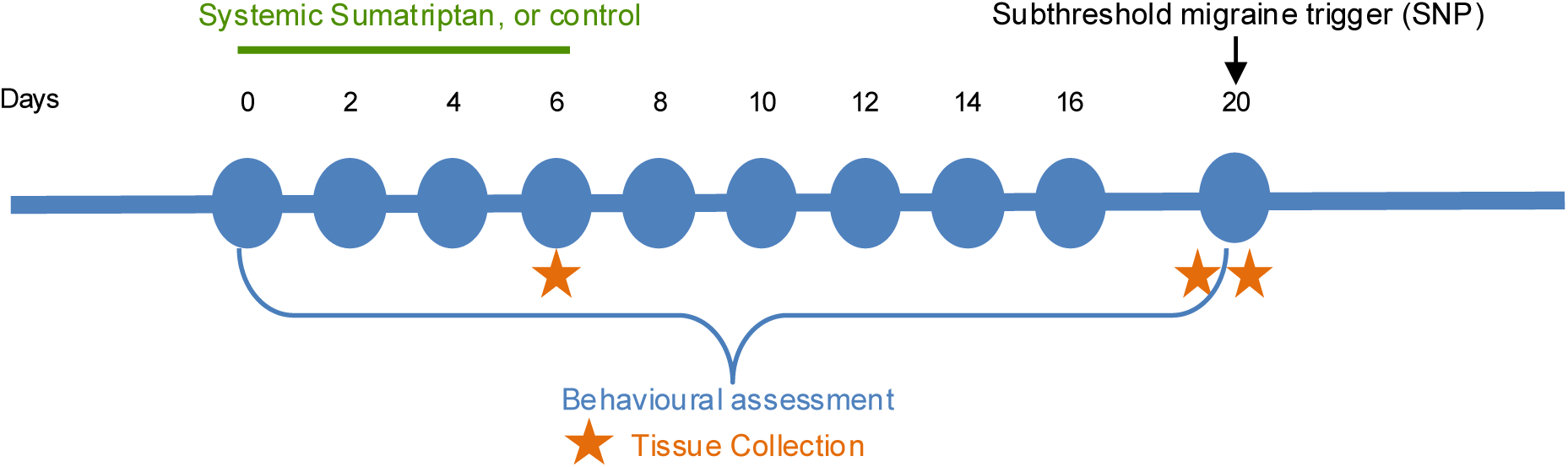
Experimental timeline. Adult male rats received continuous subcutaneous infusion of saline or sumatriptan via osmotic minipumps for 6 days. Mechanical sensitivity was assessed over 20 days. Tissues were collected at day 6 and day 20, with an additional subgroup receiving sodium nitroprusside (SNP; 3 mg/kg, i.p.) at day 20 before tissue collection.

### Evaluation of Tactile Sensitivity

At day 0 baseline withdrawal thresholds to von Frey filaments applied to the periorbital region of the face and the plantar surface of the hindpaw were determined prior to osmotic mini-pump implantation. To determine tactile sensitivity the rats were placed in cages and allowed to acclimate for 45 min. For the sensory threshold of the face or of the hindpaw, the von Frey filament was applied perpendicularly to the periorbital region of the rat’s face or to the plantar surface of the hindpaw, until it buckled slightly, and held for 3-6 s. A positive response is indicated by withdrawal of the face or hindpaw from the von Frey filament as in previous studies (23,24). Maximum filament strengths were 8 g and 15 g for the periorbital region and hindpaw, respectively. Periodic testing for cutaneous allodynia was then performed every other day starting from day 2 post minipump implantation.

### Tissue collection and preparation

Rats were transcardially perfused with 0.1 M phosphate buffered saline (PBS) followed by 4% formaldehyde/12.5% picric acid solution in 0.1 M PBS. The trigeminal ganglia (TG) and brainstem (containing trigeminal nucleus caudalis (TNC)) were removed, post fixed with 4% PAF for 4 hours, transferred overnight at 4°C to 30% sucrose, frozen and then sectioned on a cryostat. The TG sections (10 µm) were directly mounted to slides ready for immunohistochemistry, while the brainstem sections (30 µm) were processed as free-floating and then transferred to slides. Brainstem sections were cut from 5 mm caudal to 10 mm rostral to obex.

### Immunohistochemistry

TG sections were incubated overnight with primary antibody against pERK (1:600 Cell Signalling Technology), or pp38 (1:300 Cell Signalling Technology). Brainstem sections were first incubated overnight with primary antibody against pERK or pp38, and then against either GFAP (1:500 Sigma-Aldrich), Iba-1 (1:500, Abcam) or NF200 (1:500 Merck Millipore) for 24 hrs.

Secondary antibodies were donkey antirabbit/mouse immunoglobulin G (IgG) conjugated with either Cy3 or FITC (1:300 Jackson Labs) for 2 hours. Analysis of colocalisation of 2 markers was performed using dual-labelling immunofluorescence. Because the primary antibodies for the markers were raised from different species, selective secondary antibodies were employed. Brainstem sections were mounted on slides and then slides contained either TG or brainstem were cover slipped with mounting medium (Vectashield, Vector). Negative controls were included (data not shown) in all staining runs by omitting the primary antibody and replacing it with the appropriate antibody diluent; these sections showed no specific immunofluorescent signal.

### Analysis and quantification

For each marker, 6 TG and 14 TNC sections from each animal were analysed. All sections were examined using a Zeiss Axioplan 2 fluorescent microscope, and images were captured using Image Pro-Plus (version 5.1, Media Cybernetics) under identical acquisition settings within each staining experiment. In TG, pERK and pp38 neuronal profiles were quantified as the percentage of immunopositive neuronal profiles out of the total neuronal profiles counted across the whole ganglion section. In TNC, pERK and pp38 were quantified as the number of positive profiles in laminae I–II, whereas Iba-1 and GFAP were quantified by area fraction using ImageJ (version 1.50i), because these markers labelled branched cellular processes and changes in tissue coverage rather than discrete somata alone. For each section, a fixed region of interest encompassing the outer laminae of the TNC was selected using the same anatomical landmarks in all animals.

To quantify Iba-1 and GFAP staining, each image was converted using a constant pixel-intensity threshold applied identically across all sections within the same experiment. This threshold was determined empirically before analysis to distinguish specific immunofluorescent signal from background staining and was then kept unchanged for all images included in that comparison. The proportion of pixels above threshold within the defined region of interest was calculated and expressed as percentage area fraction, providing an index of overall glial marker expression within the TNC. All quantification was carried out blind to treatment group, and a selection of cell counts were repeated on different days, the totals of which were all within 5% of the original count.

### CBF measurement

All MRI measurements were made on a 7 Tesla Bruker BioSpec70/30. RF transmission utilised an 86mm (ID) 1H quadrature volume resonator and signal reception used a single channel surface coil. Rats were scanned at day 0, 6, and 20 time points under isoflurane anaesthesia (30:70 O2/N2 gas mixture). Temperature and breathing rate were monitored throughout the scan session. Baseline CBF measurement (single slice 96*96 FAIR-EPI) were performed.

### Statistical analyses

Statistical analyses were performed using SPSS Statistics (version 22.0, IBM), and graphs were prepared in GraphPad Prism 7 (version 7.0, GraphPad Software). A p value of less than 0.05 was considered statistically significant. A two-way ANOVA (with post hoc Bonferroni test) was used to compare average withdrawal thresholds in response to von Frey filaments over time between saline and sumatriptan treatment. Following SNP injection on day 20, between-group comparisons of withdrawal thresholds were performed using an independent-samples t-test.

For immunohistochemical data, saline and sumatriptan groups were compared at each experimental time point separately for each marker. Comparisons were performed using an independent-samples t-test. For rostro-caudal analyses in the TNC, each animal contributed repeated measurements across seven predefined rostro-caudal bands. These data were therefore analysed using two-way repeated-measures ANOVA, with rostro-caudal level as the within-subject factor and treatment as the between-subject factor, allowing assessment of treatment effects, spatial effects, and treatment-by-level interactions. When significant effects were detected, pairwise comparisons of estimated marginal means were performed with Bonferroni correction to identify rostro-caudal levels contributing to the effect.

## RESULTS

Periorbital and hindpaw withdrawal thresholds to von Frey were monitored every other day for a period of 20 days. As previously reported (23, 24), sumatriptan-treated animals developed significant cephalic and extracephalic mechanical hypersensitivity relative to saline controls during the infusion period, with post hoc between-group differences emerging at day 4 (Fig 2A). Following pump removal on day 6, thresholds gradually return towards baseline and had normalised by day 16, whereas saline-treated rats showed no evidence of allodynia throughout the study period (Fig 2A). On day 20, SNP challenge reduced facial and hindpaw withdrawal thresholds in previously sumatriptan-exposed rats; however, this reduction was not yet significant at the 1 hr timepoint measured (Fig 2B). Because animals were culled at 2 hours post SNP for immunohistochemistry analysis, the development of later behavioural allodynia was not assessed.

**Figure 2.**
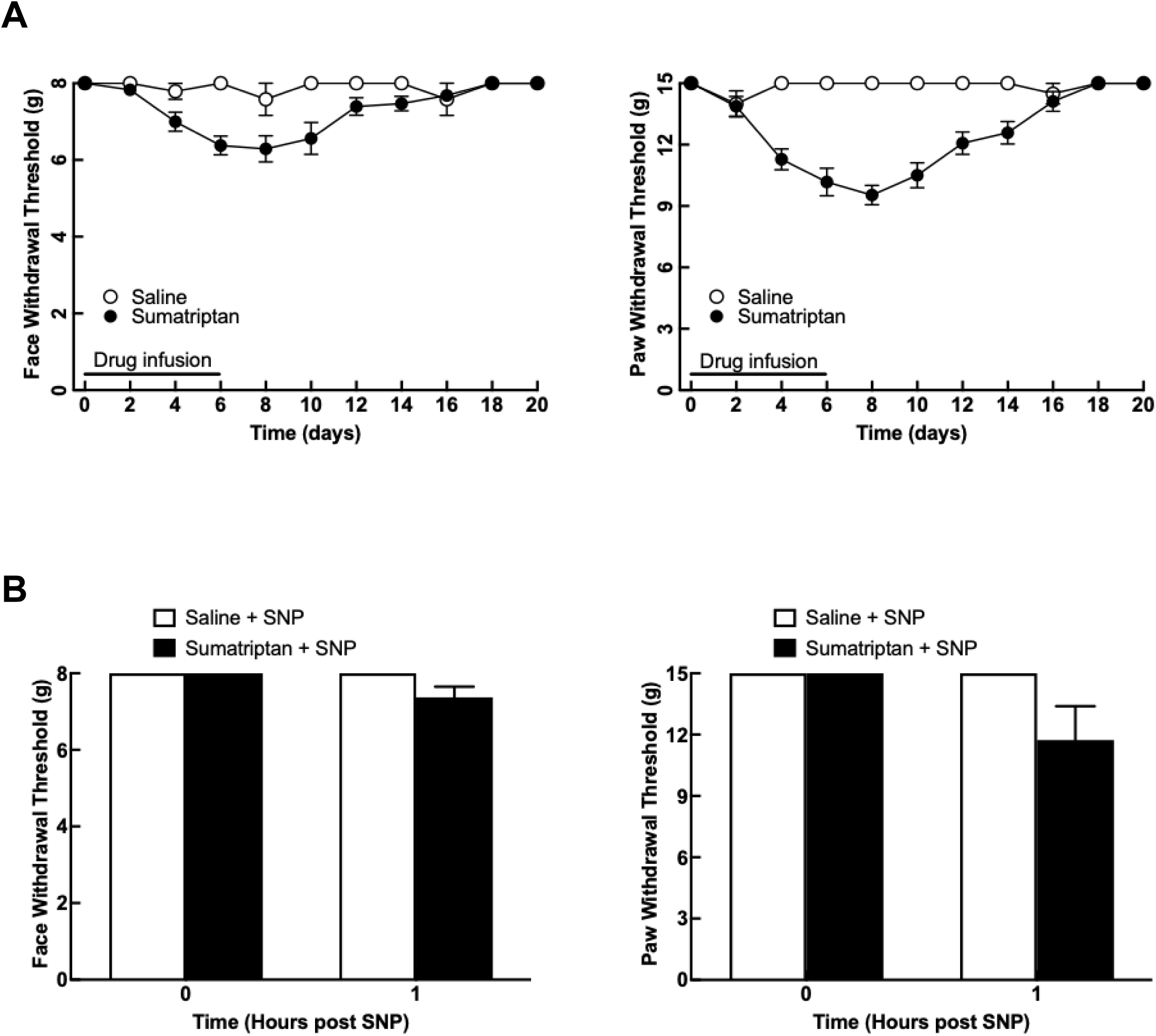
Continuous sumatriptan infusion induces reversible migraine-like mechanical hypersensitivity. (A) Periorbital and hindpaw withdrawal thresholds during and after 6 days of saline or sumatriptan infusion. Sumatriptan reduced mechanical thresholds during infusion, with recovery towards baseline after pump removal. Data were analysed by two-way ANOVA with Bonferroni post hoc testing. (B) Withdrawal thresholds at day 20 after SNP challenge. Previously sumatriptan-exposed rats showed lower thresholds than saline controls at the sampled time point, although this difference was not significant. Data are mean ± SEM.

### Increased pERK and pp38 in trigeminal ganglion following triptan treatment

Immunofluorescent labelling was used to identify trigeminal neurons expressing phosphorylated ERK (pERK) and phosphorylated p38 (pp38) in ganglion sections. Only neuronal profiles showing clear cytoplasmic or nuclear labelling and a visible nucleus were counted, while glial or non-neuronal labelling was excluded. Quantification was performed from the entire ganglion section on matched levels across animals. Sustained administration of sumatriptan for 6 days produced significant (p < 0.05) and prolonged increases in trigeminal neuronal profiles expressing pERK and pp38 (Fig. 3,4).

**Figure 3.**
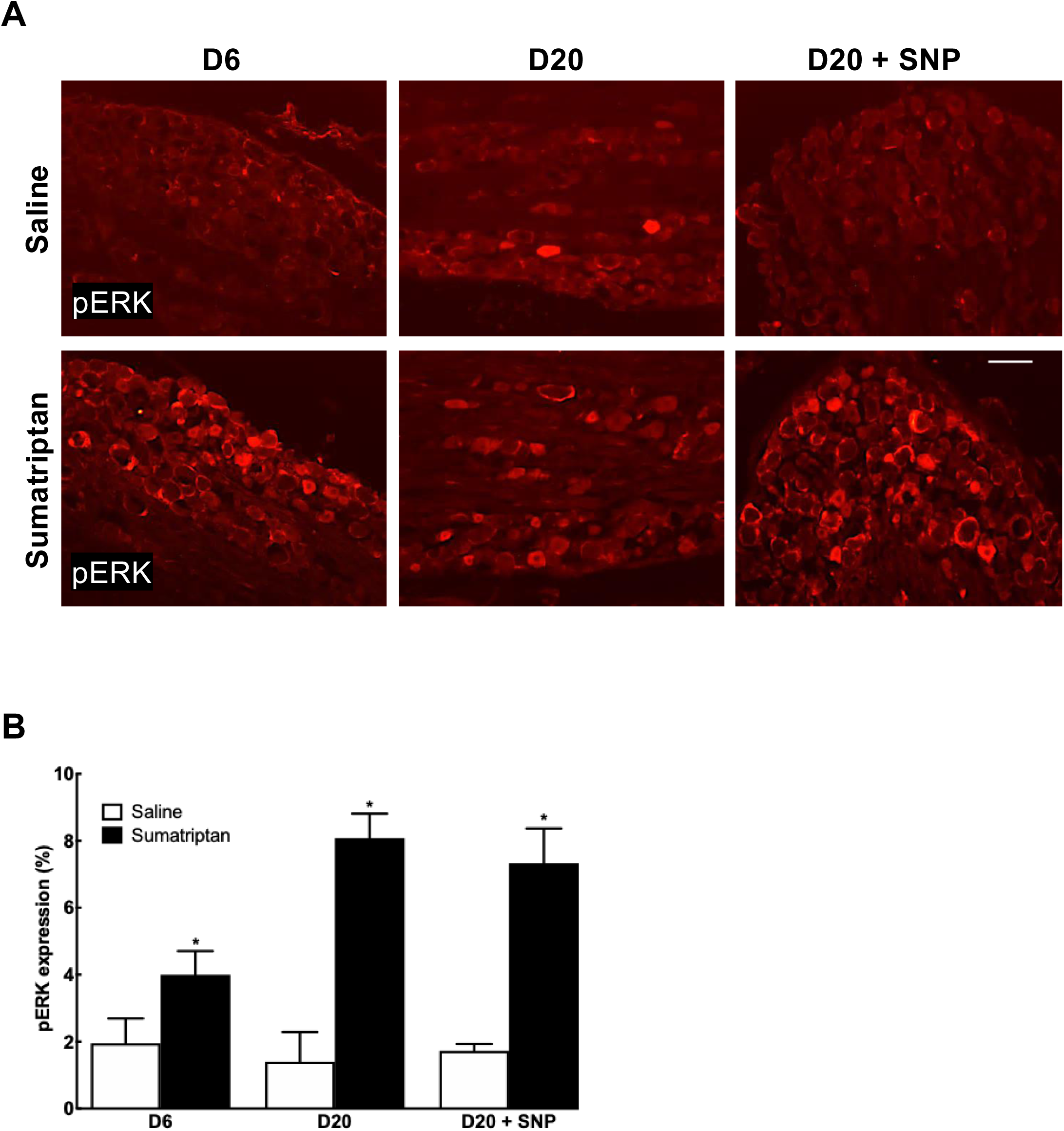
Sumatriptan exposure increases pERK expression in the trigeminal ganglion. (A) Representative immunofluorescence images of pERK labelling in trigeminal ganglion sections from saline- and sumatriptan-treated rats at day 6, day 20, and day 20 after SNP challenge. (B) Quantification of pERK-positive profiles in the trigeminal ganglion. Sumatriptan-treated rats showed increased pERK expression at day 6 and day 20 compared with saline controls (p <0.001). Values for the day 20 SNP-challenged groups are also shown. Scale bar = 100 µm; magnification 20x. Data are mean ± SEM.

**Figure 4.**
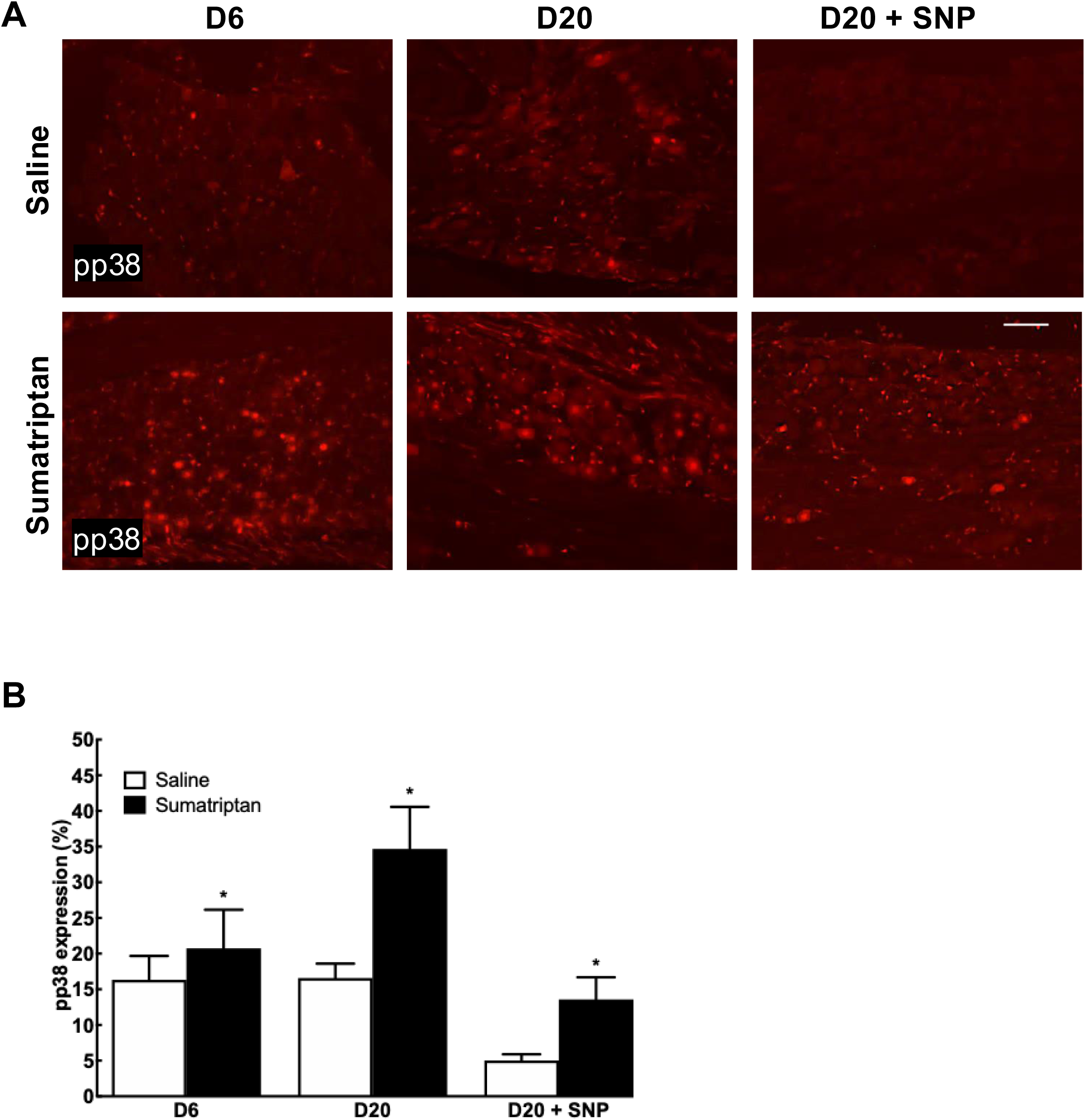
Sumatriptan exposure increases pp38 expression in the trigeminal ganglion. (A) Representative immunofluorescence images of pp38 labelling in trigeminal ganglion sections from saline- and sumatriptan-treated rats at day 6, day 20, and day 20 after SNP challenge. (B) Quantification of pp38-positive profiles in the trigeminal ganglion. Sumatriptan-treated rats showed increased pp38 expression at day 6 and day 20 compared with saline controls (p <0.001). Values for the day 20 SNP-challenged groups are also shown. Scale bar = 100 µm; magnification 20x. Data are mean ± SEM.

In the saline-treated group approximately 2% (1.9% ± 0.7 at day 6 and 1.4% ± 0.8 at day 20) of the counted trigeminal neuronal profiles were positively labelled for pERK. In contrast, sumatriptan treatment significantly increased the proportion of pERK-positive neuronal profiles to 4.0 ± 0.7% at day 6 and 8.1 ± 0.7% at day 20 (both vs. saline; p<0.001 Fig. 3A, B). After SNP challenge on day 20, pERK expression remained significantly higher in previously sumatriptan-exposed rats than in saline controls (7.33 ± 1.0% vs. 1.7 ± 0.2%; p<0.01 Fig. 3A, B).

A similar pattern was observed for pp38, although baseline labelling was higher than for pERK (Fig. 4A, B). In saline-treated rats, 16.3 ± 3.3% and 16.6 ± 2.0% of trigeminal neuronal profiles were pp38-positive at day 6 and day 20, respectively, whereas sumatriptan increased pp38-positive profiles to 20.7 ± 5.4% at day 6 and 34.7 ± 10.9% at day 20 (p<0.01 Fig. 4A, B). Following SNP administration on day 20, pp38 expression was also significantly greater in sumatriptan-pre-exposed rats than in saline-treated animals (13.6 ± 3.11% vs. 5.0 ± 0.9%; p<0.01 Fig. 4A, B).

Morphologically, pERK immunoreactivity in TG neurons appeared predominantly cytoplasmic, whereas pp38 showed a more prominent nuclear distribution. This pattern is in keeping with previous reports in pain-related tissue, in which ERK activation is often observed in the cytoplasm and dendrites, while activated p38 may accumulate more strongly in the nucleus after noxious or inflammatory stimulation. Together, these observations support the interpretation that sumatriptan treatment enhances MAPK signalling in trigeminal ganglion neurons.

### pERK and pp38 expression in satellite cells of the trigeminal ganglia

In addition to neuronal labelling, both pERK and pp38 immunoreactivity were also observed in non-neuronal cells within the trigeminal ganglion, consistent with satellite glial cells. To distinguish neuronal from non-neuronal expression, sections from day 6 saline- and sumatriptan-treated animals were dual-labelled for either pERK or pp38 together with the neuronal marker NeuN.

In sumatriptan-treated animals, much of the increased pERK signal was associated with NeuN-positive neuronal cell bodies, indicating that the enhanced pERK expression shown in Figure 3 primarily occurred in primary afferent neurons (Fig. 5). However, pERK immunoreactivity was not restricted to neurons, and labelled non-neuronal cells were also present in both saline- and sumatriptan-treated ganglia. A similar pattern was observed for pp38, with neuronal localisation accounting for much of the increased signal in sumatriptan-treated animals, while additional non-neuronal labelling was also evident, although representative images are not shown.

**Figure 5.**
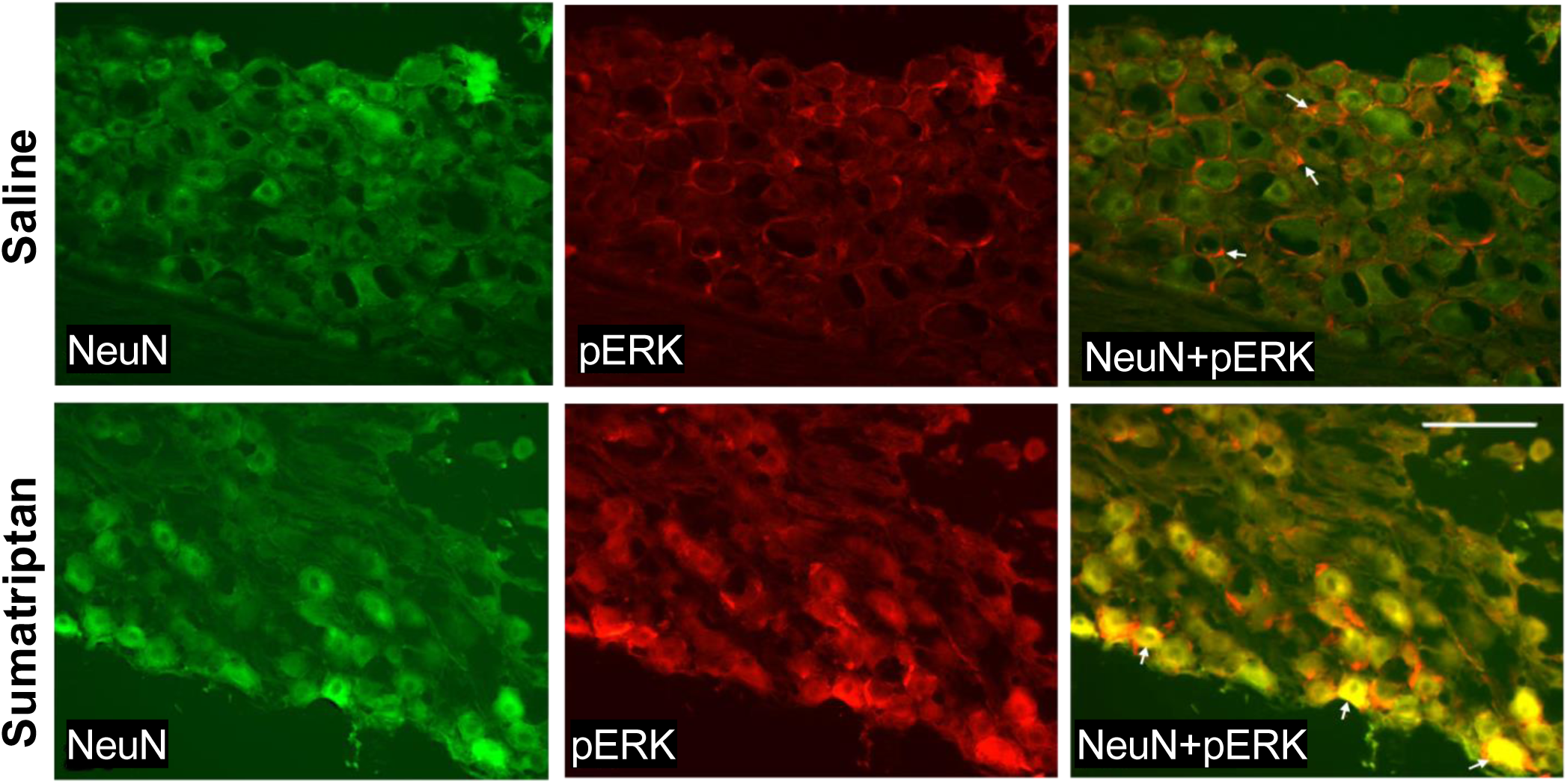
pERK expression in neuronal and non-neuronal cells of the trigeminal ganglion. Representative trigeminal ganglion sections from day 6 saline- and sumatriptan-treated rats dual-labelled for pERK and the neuronal marker NeuN. Increased pERK signal in sumatriptan-treated animals is largely associated with NeuN-positive neuronal profiles, although non-neuronal pERK labelling is also present.

### Increased pERK and pp38 in trigeminal nucleus caudalis following triptan treatment

Immunofluorescent labelling was used to quantify pERK- and pp38-positive cells in the outer laminae of the trigeminal nucleus caudalis (TNC). Expression was assessed at day 6, day 20, and 2 h after SNP challenge on day 20.

pERK expression in the TNC was significantly increased after 6 days of sumatriptan infusion compared with saline treatment (5.4 ± 0.6 vs. 3.0 ± 0.5; p<0.001 Fig. 6A, B). This increase persisted after discontinuation of sumatriptan, such that pERK expression remained significantly higher in the sumatriptan group than in saline controls at day 20 (31.8 ± 3.2 vs. 4.4 ± 1.1; p<0.001 Fig. 6A, B). Following SNP administration on day 20, pERK expression increased in both treatment groups relative to day 20 without SNP (p<0.01). In saline-treated rats, pERK increased from approximately 4.0 cells at day 20 to 27.9 ± 2.3 cells 2 h after SNP, whereas in previously sumatriptan-exposed rats pERK expression reached 10.7 ± 0.9 cells (Fig. 6A, B). Thus, although pERK remained elevated in the sumatriptan group relative to saline controls without SNP (p<0.01), the response in the sumatriptan plus SNP group was lower than that observed in the saline plus SNP group (Fig. 6A, B).

**Figure 6.**
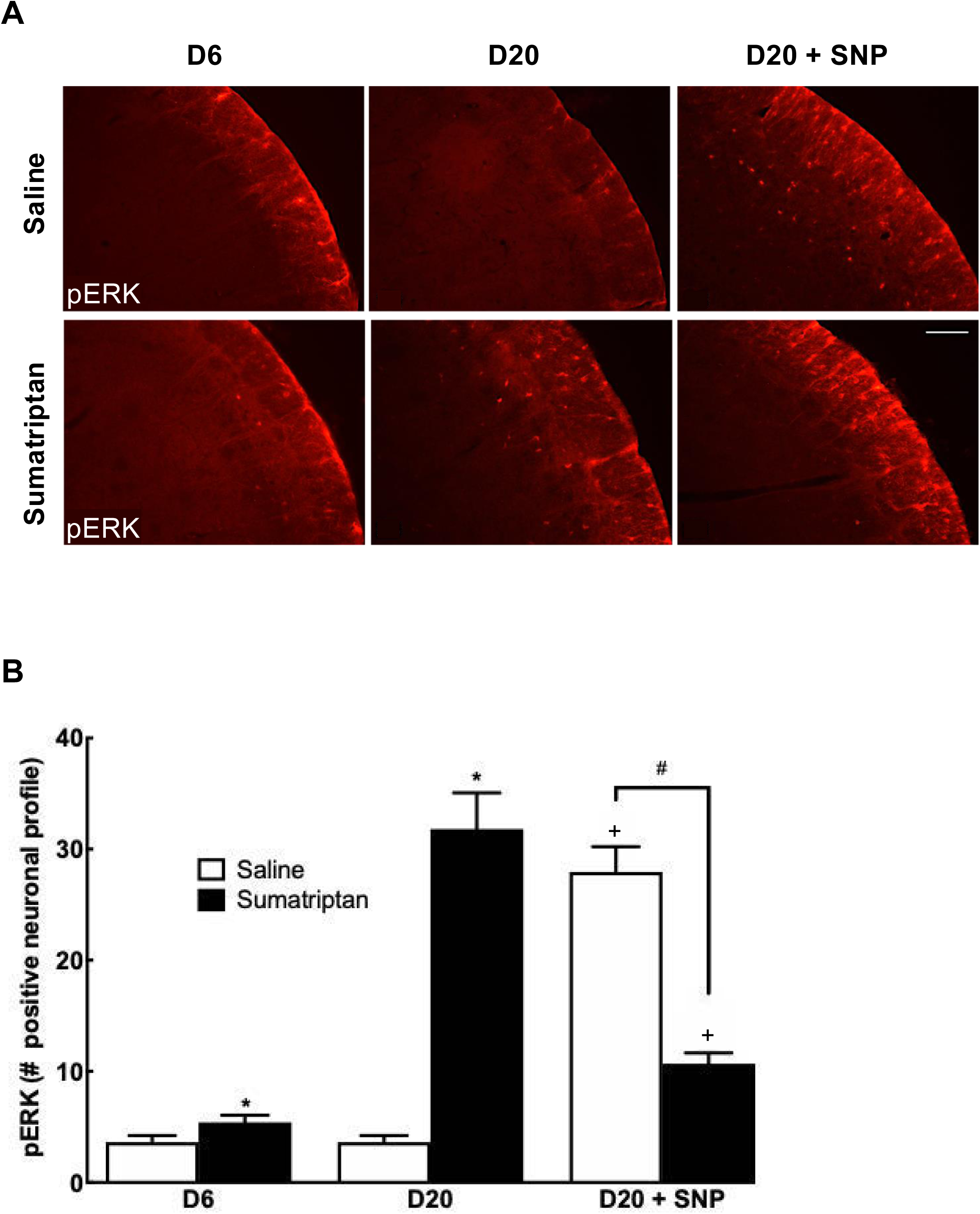
Sumatriptan exposure increases pERK expression in the trigeminal nucleus caudalis. (A) Representative immunofluorescence images of pERK labelling in the outer laminae (I-II) of the trigeminal nucleus caudalis from saline- and sumatriptan-treated rats at day 6, day 20, and day 20 after SNP challenge. (B) Quantification of pERK-positive profiles in the trigeminal nucleus caudalis. Sumatriptan-treated rats showed increased pERK expression at day 6 (*p <0.05) and day 20 (*p <0.001) compared with saline controls. SNP challenge increased pERK expression in both groups, with a larger response in saline-treated rats (+p <0.001) compared to saline control at day 6. Significant difference was found between saline and sumatriptan treated rats plus SNP (# p <0.001). Scale bar = 100 µm; magnification 20x. Data are mean ± SEM.

A similar overall pattern was observed for pp38 (Fig. 7A, B). After 6 days of sumatriptan infusion, pp38 expression in the TNC was significantly greater than in saline-treated rats (115.0 ± 13.3 vs. 45.4 ± 5.4; p<0.001 Fig. 7A, B). Elevated pp38 expression persisted at day 20 after drug discontinuation, remaining higher in the sumatriptan group than in saline controls (96.5 ± 10.8 vs. 11.1 ± 1.7; p<0.001 Fig. 7A, B). SNP challenge on day 20 increased pp38 expression in both groups compared with saline day 20 values. In saline-treated rats, pp38 increased to 81.1 ± 4.2, whereas in previously sumatriptan-exposed rats pp38 reached 75.0 ± 5.6 (p<0.01) Fig. 7A, B). Unlike day 6 and day 20 without SNP, pp38 expression did not differ clearly between the saline plus SNP and sumatriptan plus SNP groups (p=0.5 Fig. 7A, B).

**Figure 7.**
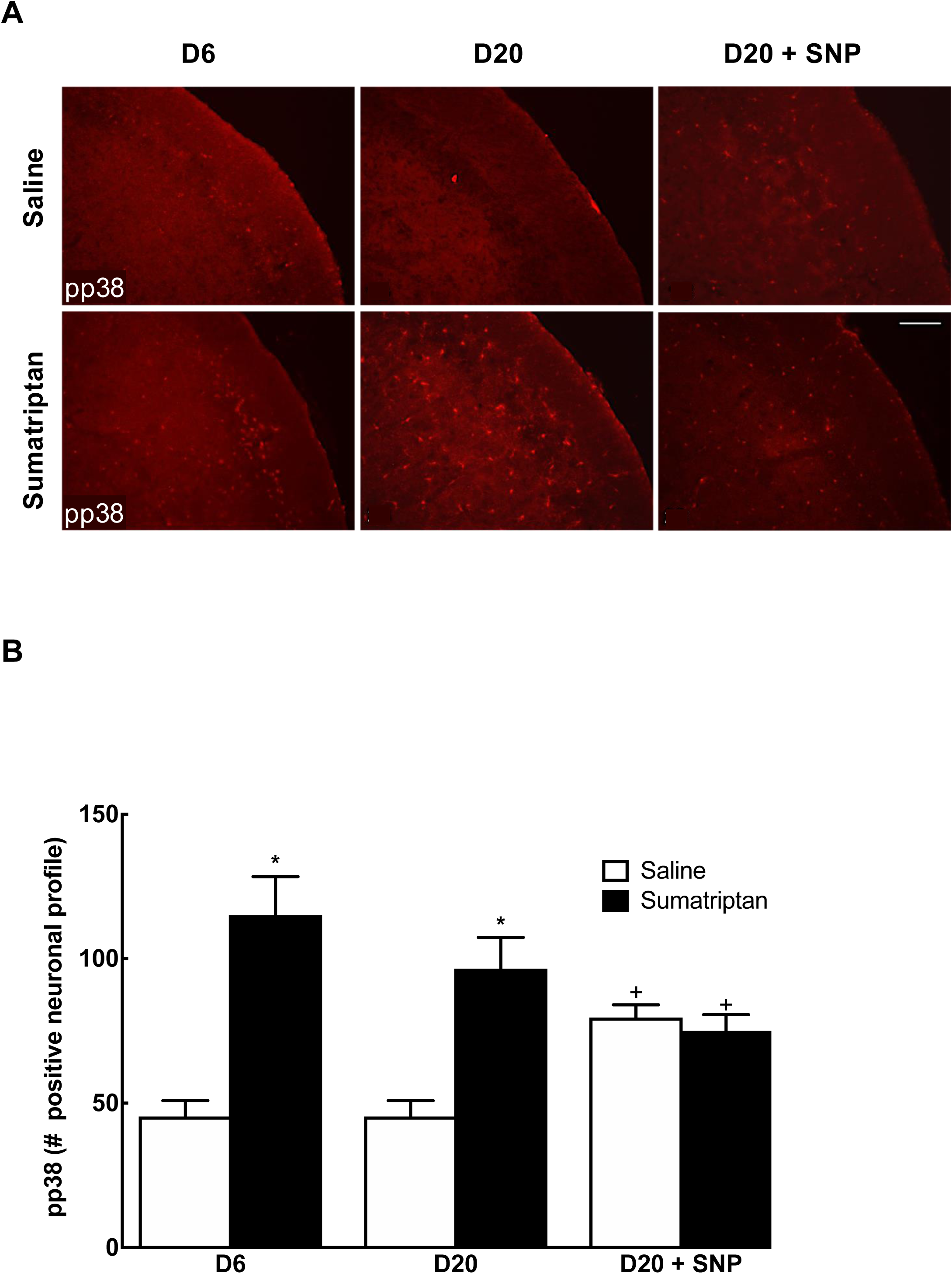
Sumatriptan exposure increases pp38 expression in the trigeminal nucleus caudalis. (A) Representative immunofluorescence images of pp38 labelling in the outer laminae (I-II) of the trigeminal nucleus caudalis from saline- and sumatriptan-treated rats at day 6, day 20, and day 20 after SNP challenge. (B) Quantification of pp38-positive profiles in the trigeminal nucleus caudalis. Sumatriptan-treated rats showed increased pp38 expression at day 6 and day 20 compared with saline controls (*p <0.001). SNP challenge increased pp38 expression in both groups (+p <0.001), and no clear difference was observed between saline and sumatriptan rats after SNP challenge (p = 0.26). Scale bar = 100 µm; magnification 20x. Data are mean ± SEM.

### Increased activation of microglial and astrocytic in the trigeminal nucleus caudalis following triptan treatment

Immunofluorescent labelling for Iba-1 and GFAP was used to assess microglial and astrocytic activation, respectively, in the trigeminal nucleus caudalis (TNC). Expression was quantified as the percentage area of positive labelling within a standardised region of the outer TNC at day 6, day 20, and 2 h after SNP challenge on day 20.

Iba-1 expression was not significantly altered after 6 days of sumatriptan infusion compared with saline treatment (2.2 ± 0.2% vs. 1.8 ± 0.2%; Fig. 8A, B). However, by day 20, after discontinuation of treatment, Iba-1 expression was significantly increased in the sumatriptan group relative to saline controls (4.2 ± 0.4% vs. 1.9 ± 0.4%; Fig. 8A, B). SNP challenge further increased Iba-1 expression in both groups, reaching 4.9 ± 0.3% in saline-treated rats and 6.0 ± 0.4% in previously sumatriptan-exposed rats (Fig. 8A, B). Thus, microglial activation was most pronounced after treatment withdrawal and was further enhanced by SNP in sumatriptan-pre-exposed animals.

**Figure 8.**
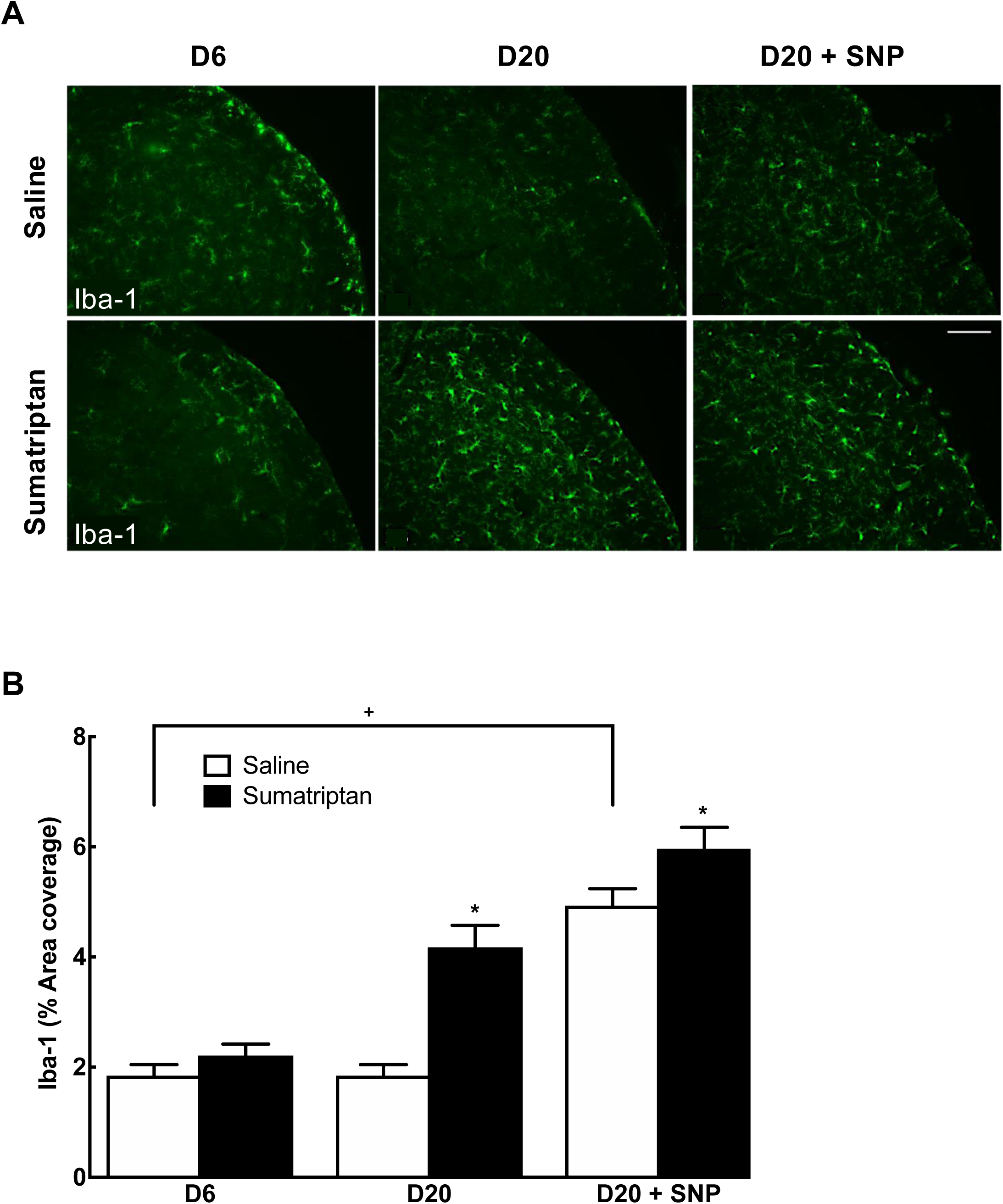
Microglial activation in the trigeminal nucleus caudalis following sumatriptan exposure. (A) Representative immunofluorescence images of Iba-1 labelling in the trigeminal nucleus caudalis from saline-and sumatriptan-treated rats at day 6, day 20, and day 20 after SNP challenge. (B) Quantification of Iba-1-positive area. No significant difference was observed at day 6, whereas microglial labelling was increased at day 20 in sumatriptan-treated rats compared with saline controls (*p <0.001). SNP challenge further increased Iba-1 expression in both groups (+p <0.001), with the greatest values in previously sumatriptan-exposed rats (*p=0.001). Representative images also show morphological changes consistent with glial activation. Scale bar = 100 µm; magnification 20x. Data are mean ± SEM.

GFAP expression showed a similar but more delayed profile (Fig. 9A, B). At day 6, GFAP labelling was not significantly different between sumatriptan- and saline-treated rats (2.0 ± 0.3% vs. 1.6 ± 0.3%; Fig. 9A, B). At day 20, GFAP expression was significantly increased in the sumatriptan group compared with saline controls (2.4 ± 0.2% vs. 1.7 ± 0.2%; Fig. 9A, B). Following SNP challenge, GFAP expression increased further in both groups, reaching 3.9 ± 0.3% in saline-treated rats and 4.8 ± 0.4% in previously sumatriptan-exposed rats (Fig. 9A, B). These findings indicate that both microglial and astrocytic activation persist after cessation of sumatriptan treatment and are further augmented by SNP challenge.

**Figure 9.**
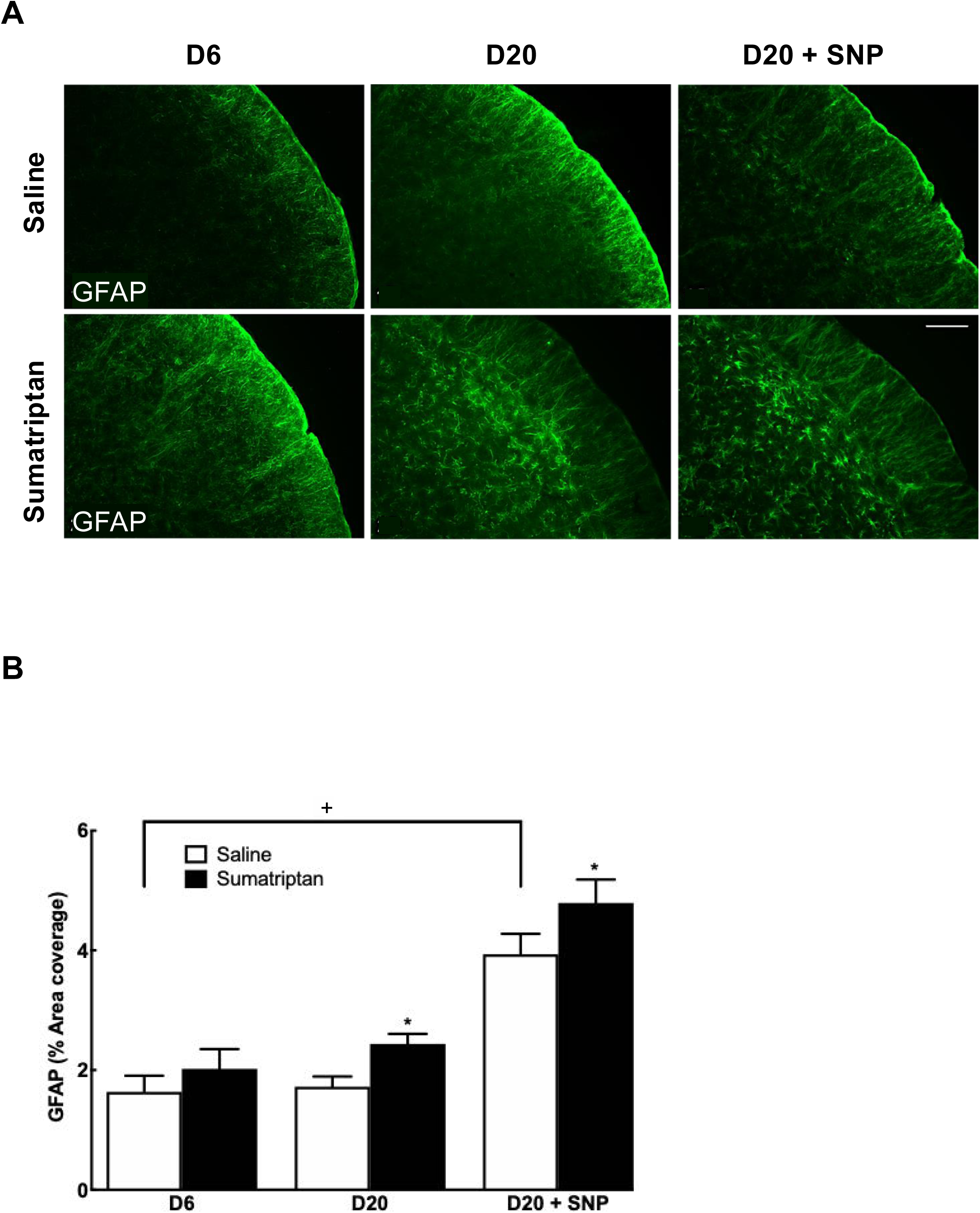
Astrocytic activation in the trigeminal nucleus caudalis following sumatriptan exposure. (A) Representative immunofluorescence images of GFAP labelling in the trigeminal nucleus caudalis from saline-and sumatriptan-treated rats at day 6, day 20, and day 20 after SNP challenge. (B) Quantification of GFAP-positive area. No significant difference was observed at day 6, whereas astrocytic labelling was increased at day 20 in sumatriptan-treated rats compared with saline controls (*p <0.001). SNP challenge further increased GFAP expression in both groups (+p <0.001), with the greatest values in previously sumatriptan-exposed rats (*p <0.001). Representative images also show morphological changes consistent with glial activation. Scale bar = 100 µm; magnification 20x. Data are mean ± SEM.

In addition to the quantitative changes, morphological alterations suggestive of glial activation were observed at day 20 and after SNP challenge, including hypertrophy and reduced ramification, particularly in the sumatriptan-treated group (Figs. 8A and 9A). Because these observations are based on representative images, they are best described qualitatively rather than used as a primary quantitative outcome.

### Rostral-caudal distribution of biomarker expression in the TNC

To examine the spatial distribution of biomarker expression within the trigeminal nucleus caudalis (TNC), sections from day 6 animals were grouped into seven predefined rostro-caudal bands according to their distance from the obex, and mean expression values were calculated for each band for each animal. Because each animal contributed measurements across multiple rostro-caudal levels, these data were analysed using two-way repeated-measures ANOVA, with rostro-caudal level as the within-subject factor and treatment as the between-subject factor. This approach accounts for the non-independence of measurement obtained from multiple rostro-caudal levels within the same animal.

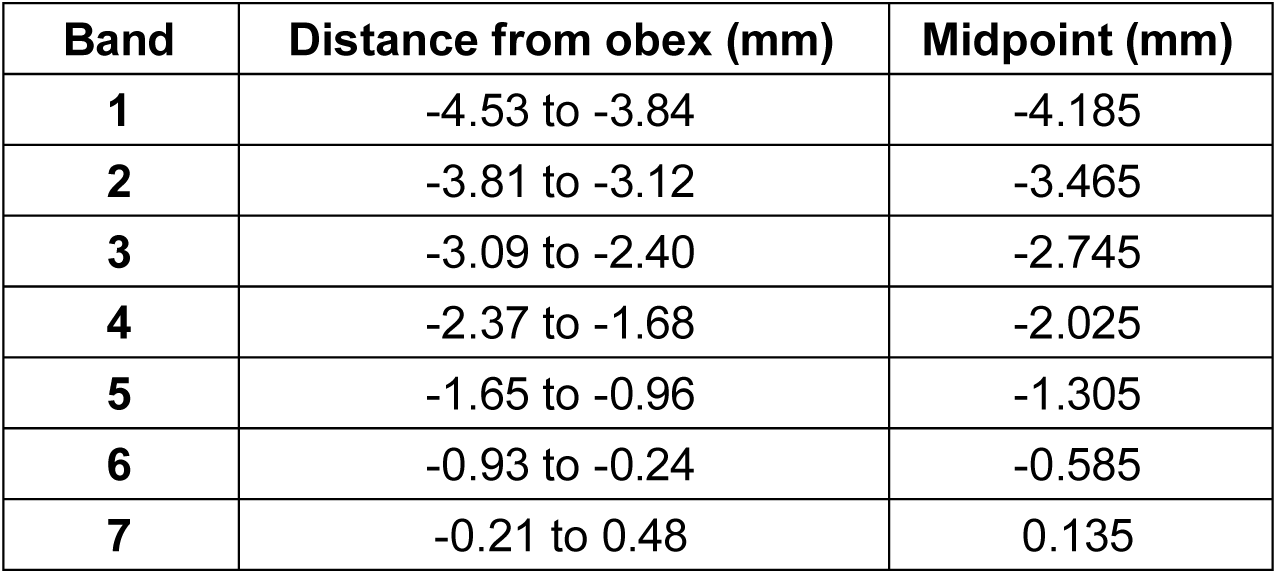

| Band | Distance from obex (mm) | Midpoint (mm) |
| --- | --- | --- |
| 1 | -4.53 to -3.84 | -4.185 |
| 2 | -3.81 to -3.12 | -3.465 |
| 3 | -3.09 to -2.40 | -2.745 |
| 4 | -2.37 to -1.68 | -2.025 |
| 5 | -1.65 to -0.96 | -1.305 |
| 6 | -0.93 to -0.24 | -0.585 |
| 7 | -0.21 to 0.48 | 0.135 |

Both pp38 and pERK showed regionally varying expression across the rostro-caudal axis, with the greatest separation between saline- and sumatriptan-treated animals occurring in the caudal TNC, around 2 mm caudal to the obex (Fig. 10A, B). By contrast, Iba-1 and GFAP showed a more general decline towards the obex and less obvious treatment-related separation across bands (Fig. 10C, D). The repeated-measures ANOVA identified a significant main effect of treatment for pERK and pp38, but not for Iba-1 or GFAP.

**Figure 10.**
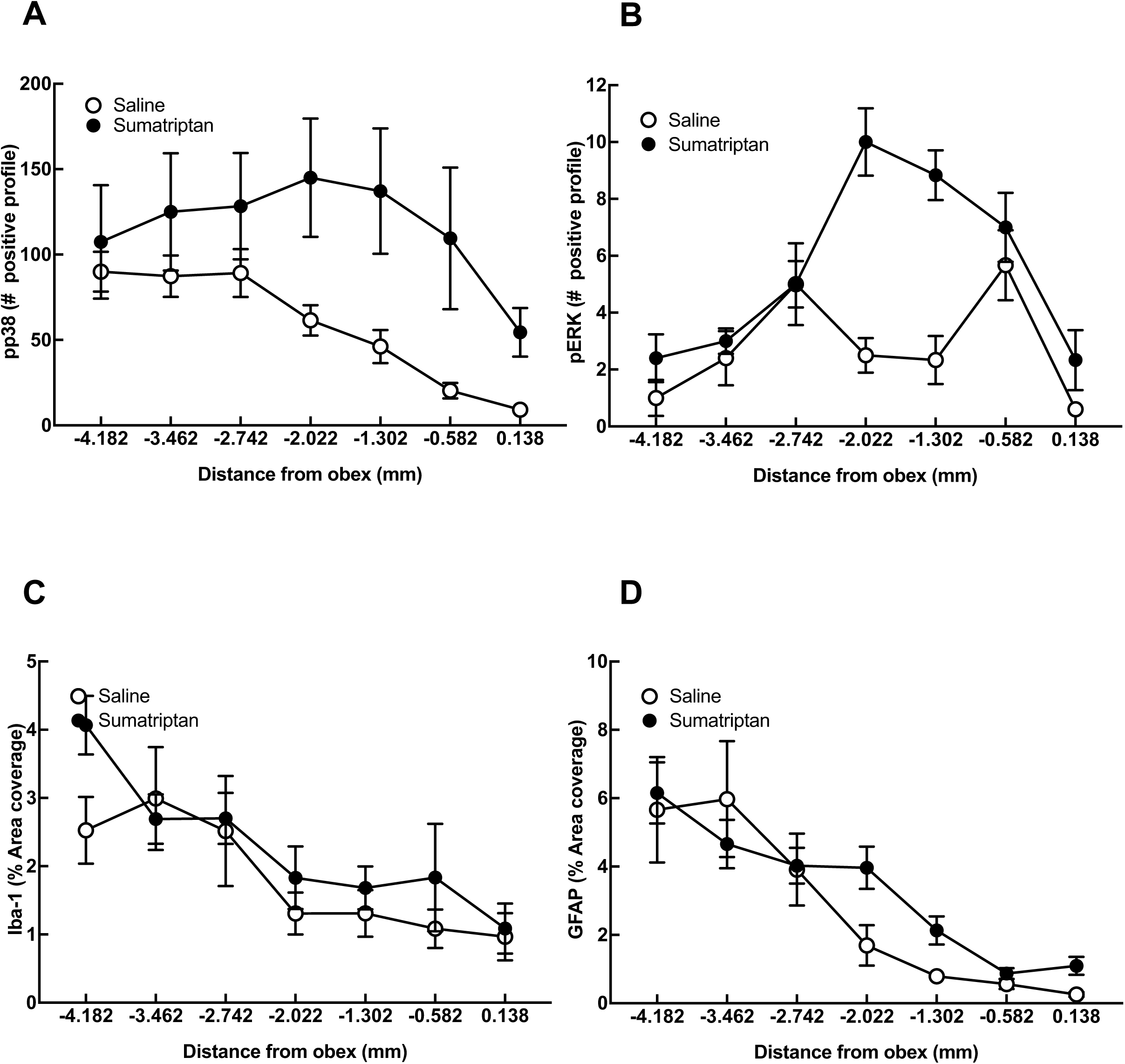
Rostro-caudal distribution of biomarker expression in the trigeminal nucleus caudalis at day 6. Expression is shown by distance from the obex for pp38 (A), pERK (B), Iba-1 (C), and GFAP (D) in saline- and sumatriptan-treated rats. pERK and pp38 showed the greatest separation between treatment groups in the caudal trigeminal nucleus caudalis, whereas Iba-1 and GFAP showed a more general decline towards the obex. Data are mean ± SEM.

### Biomarker colocalisation in the trigeminal nucleus caudalis

Tissues from all treatment groups were dual-labelled for pERK and pp38 with NeuN, Iba-1 and GFAP in the trigeminal nucleus caudalis (TNC) (Figs. 11–13 and Supplementary Figs. S1–S3). These images were used to assess the apparent cellular association of pERK and pp38 immunoreactivity across treatment groups. At day 6 following sumatriptan infusion, pERK immunoreactivity was predominantly associated with NeuN-positive neuronal profiles (Fig. 11). By day 20, with or without SNP challenge, neuronal pERK labelling appeared less prominent, while greater apparent association with glial markers was observed in the supplementary dual-labelling images. Dual-labelling of pERK with GFAP also showed biomarker signal in regions of increased astrocytic labelling, particularly at day 20 and after SNP challenge (Fig. 12). Similarly, pp38 immunoreactivity at day 6 was mainly associated with NeuN-positive neuronal profiles, particularly in sumatriptan-treated animals (Fig. 13), whereas at day 20 pp38 showed greater apparent association with Iba-1-positive microglial profiles and less evident neuronal labelling in the sumatriptan group (Supplementary Fig. S1). Dual-labelling of pERK with Iba-1 and of pp38 with GFAP further showed biomarker signal in regions of increased glial labelling at later time points (Supplementary Figs. S2 and S3). Because these observations are qualitative, they should be interpreted cautiously. Nevertheless, taken together they support a time-dependent change from predominantly neuronal biomarker association at day 6 to greater apparent glial association at day 20.

**Figure 11.**
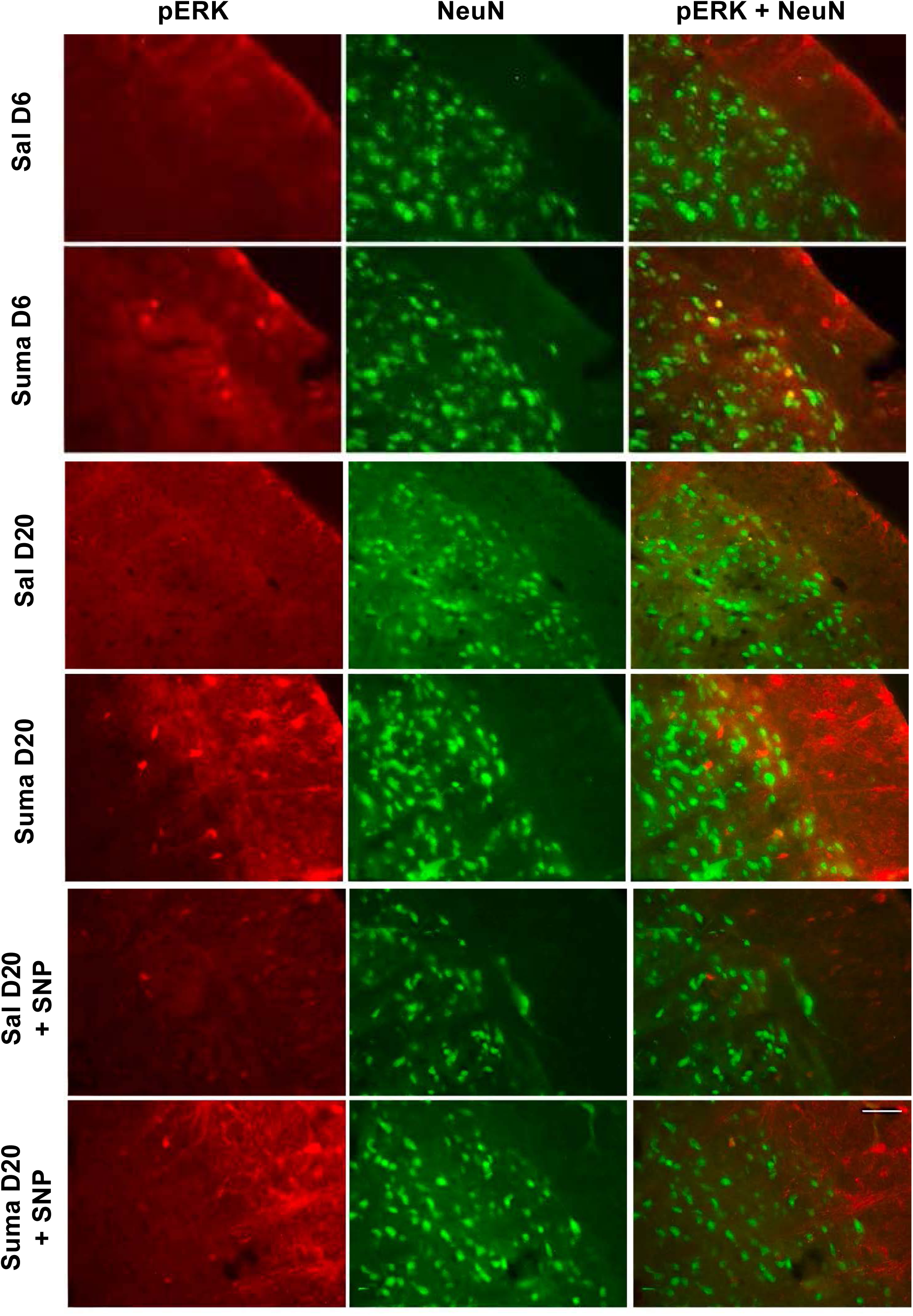
Representative dual-labelling of pERK and NeuN in the trigeminal nucleus caudalis. Images show saline- and sumatriptan-treated rats at day 6, day 20, and day 20 after SNP challenge. At day 6, pERK signal is predominantly associated with NeuN-positive neuronal profiles.

**Figure 12.**
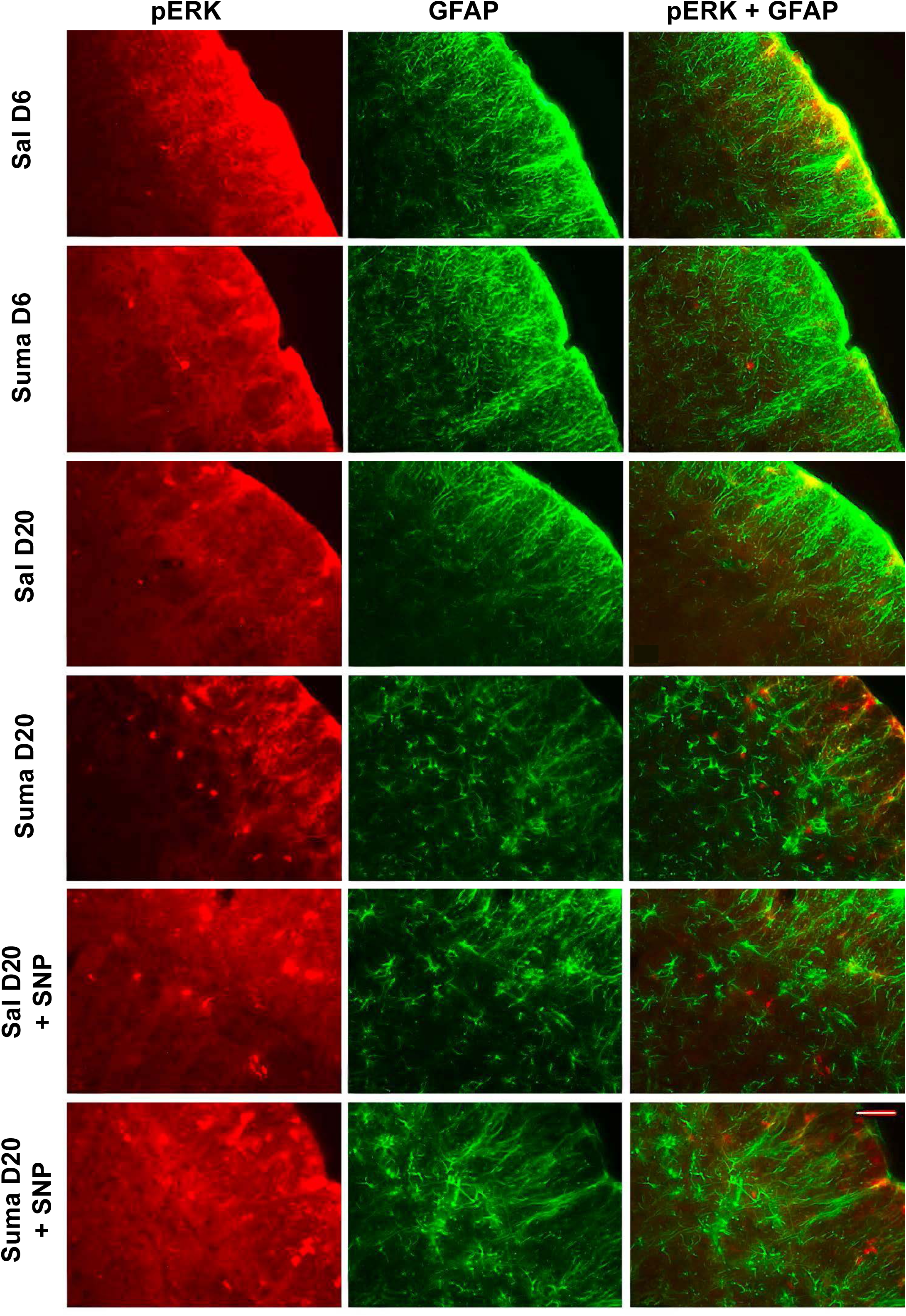
Representative dual-labelling of pERK and GFAP in the trigeminal nucleus caudalis. Images show saline- and sumatriptan-treated rats at day 6, day 20, and day 20 after SNP challenge. These panels qualitatively illustrate pERK signal in regions of increased astrocytic labelling.

**Figure 13.**
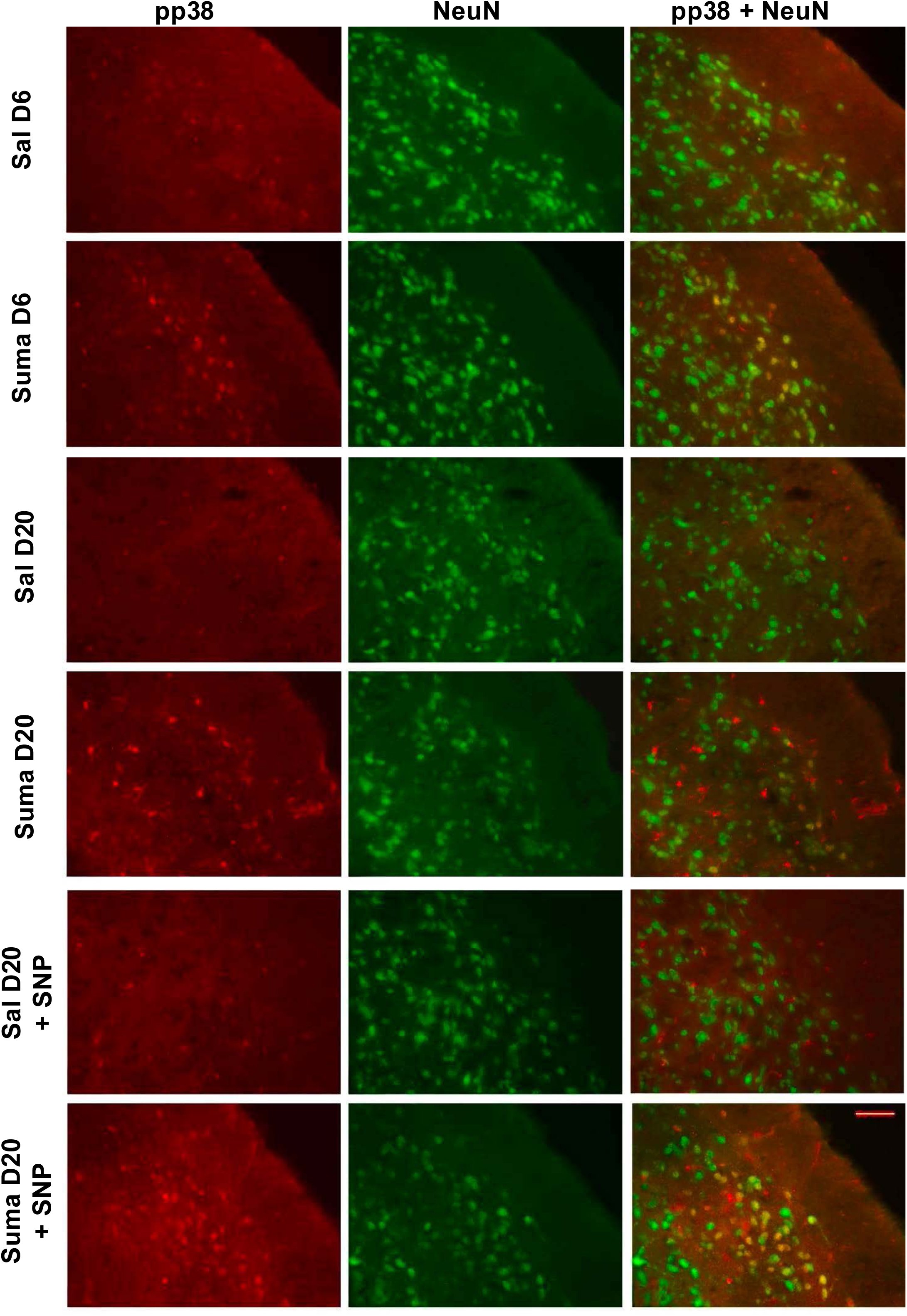
Representative dual-labelling of pp38 and NeuN in the trigeminal nucleus caudalis. Images show saline- and sumatriptan-treated rats at day 6, day 20, and day 20 after SNP challenge. pp38 signal shows neuronal association, particularly at day 6.

### Brain activity alterations in sumatriptan-treated rats

MRI assessment showed that continuous sumatriptan administration reduced cerebral blood flow (CBF) in grey matter structures compared with saline-treated controls (Fig. 14A, B). This reduction was evident after 6 days of treatment and remained present at day 20, 14 days after discontinuation of sumatriptan, indicating that altered brain perfusion persisted beyond the period of overt allodynia. These data suggest that repeated sumatriptan exposure is associated with longer-lasting changes in brain perfusion.

**Figure 14.**
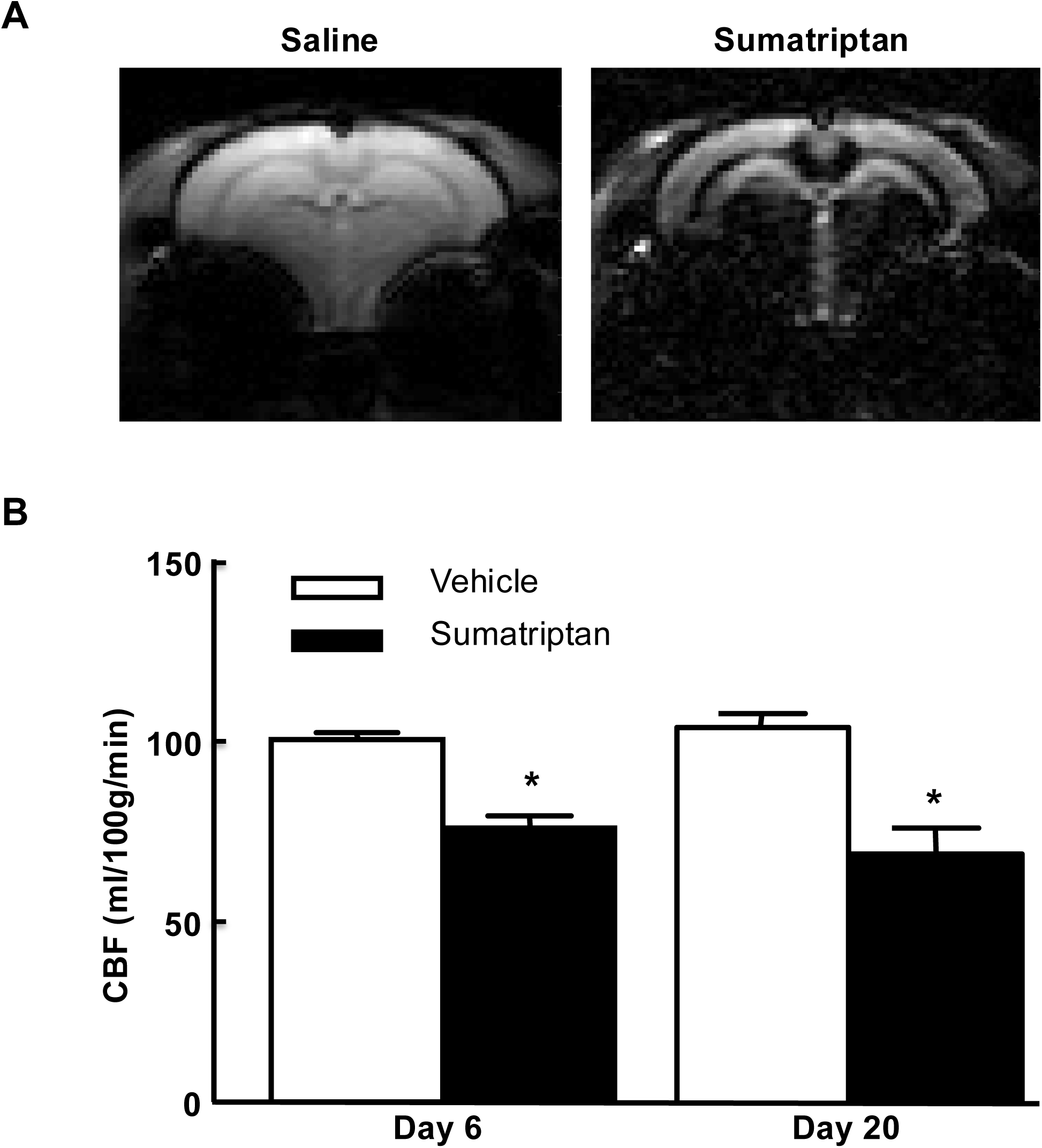
Continuous sumatriptan exposure is associated with persistent reduction in cerebral blood flow. (A) Representative FAIR-EPI MRI images acquired after saline or sumatriptan treatment. (B) Quantification of cerebral blood flow showed reduced CBF in sumatriptan-treated rats compared with saline controls (*p <0.01), and this reduction remained evident after withdrawal of the drug. Data are mean ± SEM.

## DISCUSSION

Studying migraine pain preclinically remains challenging because migraine-like pain occurs in the absence of overt tissue injury and is characterised by recurrent, intermittent episodes. We have previously shown that continuous sumatriptan administration in rats reproduces several features relevant to migraine and medication overuse headache. In the present study, we reasoned that the increased sensitivity induced by persistent triptan exposure reflects peripheral and central sensitisation within the trigeminovascular system (TGVS). Our data show that continuous sumatriptan exposure increases pERK and pp38 expression within the TGVS. Moreover, in rats previously exposed to sumatriptan, we found increased glial activation within the trigeminal nucleus caudalis (TNC), particularly after drug withdrawal, which may contribute to latent sensitisation. We also found that these changes within the TGVS were accompanied by a persistent decrease in cerebral blood flow (CBF), suggesting that repeated triptan exposure induces longer-lasting adaptations beyond the period of overt allodynia.

### Peripheral and central sensitisation in response to sumatriptan infusion

Our finding of increased MAP kinase expression within the TGVS is consistent with previous reports. pERK expression has been shown to increase in the trigeminal ganglion (TG) in response to cortical spreading depression and dural inflammation induced by capsaicin (38), and in both the TG and TNC in response to nitroglycerin infusion (39). In contrast, pp38 expression in the TG and TNC has been less well characterised in migraine-relevant models, although findings from other models of neuropathic pain support its involvement in sensitisation.

An interesting observation in the present study was that triggering a migraine-like state with SNP resulted in a substantial increase in biomarker expression in saline-treated animals as well, particularly for pERK, which was expressed at a higher level in this group than in the sumatriptan-pretreated group. The lower pERK response observed in previously sumatriptan-exposed rats may reflect altered responsivity of this pathway after prior sustained activation. However, this interpretation remains speculative and was not directly tested in the present study.

### Somatotopical expression of pERK and pp38 in the TNC in sumatriptan-treated rats

The neurons of the dorsal horn of the spinal cord are organised by function into laminae first described by Rexed (40). The dorsal horn is continuous with the TNC, and the outermost laminae (I-II) of the TNC have been identified as the main site of termination of peripheral nociceptors and the second-order afferents with which they synapse. In particular, this region is thought to represent the termination zone of C-afferent fibres (41). Nerve fibres of this type within the meninges are involved in the release of inflammatory mediators during neurogenic inflammation, and their sensitisation contributes to the cutaneous allodynia observed in migraine, as the wide dynamic range neurons with which they synapse in the TNC receive convergent input from other fibre types, including mechanoreceptive A-β fibres (42,43). The biomarker expression of both neuronal and non-neuronal origin observed in this study was concentrated in laminae I and II of the TNC, suggesting involvement of neurons receiving nociceptive input relevant to migraine pain pathways. The restriction of biomarker changes to the ophthalmic representation within the TNC further supports the view that systemic sumatriptan induces sensitisation specifically in migraine-relevant craniofacial nociceptive pathways.

Secondary afferent neurons in the TNC are organised somatotopically along the rostro-caudal axis according to the peripheral territory from which they receive sensory input (26,40–43). In the present study, the enhanced expression of pERK and pp38 observed in day 6 sumatriptan-treated rats was most prominent approximately 2 mm caudal to the obex. This region corresponds to the ophthalmic representation of the trigeminal nerve (V1), which also includes dural afferents relevant to migraine pathophysiology (26,43). 5-HT receptors, particularly 5-HT1B, are known to be expressed in these afferents, and if this is a relevant site of action of sumatriptan in MOH, it would be consistent for these biomarkers to be most prominent in the somatotopically relevant region, as observed here. Although the marked difference in expression between saline- and sumatriptan-treated animals at this level could potentially be viewed as contributing disproportionately to the overall treatment effect, repeating the analysis with band 4 (-2.37 to -1.68 mm) excluded still showed increased expression in the sumatriptan group compared with the saline group.

GFAP and Iba-1 expression followed a different pattern from that of pERK and pp38, with a general decline in expression moving rostrally towards the obex. As the difference between saline- and sumatriptan-treated animals was less marked for these glial markers, it is unlikely that the rostro-caudal distribution of glial activation is as strongly somatotopically related to migraine-relevant nociceptive pathways as that of pERK and pp38.

### Glial activation and maintenance of sensitisation

An important finding of this study was the delayed increase in Iba-1 and GFAP expression in the TNC after discontinuation of sumatriptan treatment. Although glial marker expression was not markedly altered at day 6, both microglial and astrocytic labelling were increased at day 20, and this response was further enhanced by SNP challenge. These findings suggest that glial activation is more closely associated with maintenance or reactivation of a sensitised state than with its initial induction in this model.

Microglia and astrocytes are increasingly recognised as active participants in pain-related plasticity rather than simply supportive cells. Microglia can be activated by neuronal signals, including chemokines, purines such as ATP, and glutamate (44–46). Astrocytic activation is often considered more delayed and may represent a more prolonged component of sensitisation, which is broadly consistent with the present findings. As with microglia, bidirectional communication between neurons and astrocytes may influence the function of both cell types. Activated microglia release a range of mediators, including interleukins, TNF-α and brain-derived neurotrophic factor (BDNF), all of which have been implicated in sensitisation of nociceptive neurons (44–46). One possible route for microglia-induced neuronal activation involves ATP-sensitive P2X receptors, particularly P2RX4. Tsuda et al. (47) reported that intraspinal administration of exogenous microglia with activated P2RX4 elicited tactile allodynia in the rat. BDNF has also been implicated in central sensitisation, and its application to spinal lamina I neurons produces a depolarising effect similar to that observed following microglial activation by ATP through P2X receptors (48,49). In addition, Long and colleagues (2020) (50) showed in a mouse model of chronic migraine that P2X4R-mediated BDNF release contributes to central sensitisation and neuronal hyperexcitability in the TNC. While the present data do not establish a specific glial signalling pathway in this model, they are consistent with the possibility that persistent glial activation contributes to maintenance of the sensitised state following repeated sumatriptan exposure.

### Sumatriptan-induced latent sensitisation and MOH: a possible glial contribution

An interesting observation from the dual-labelling experiments was that TNC expression of both pERK and pp38 at day 20 appeared less prominently associated with neuronal profiles and showed greater apparent association with glial markers than at day 6, when expression was predominantly neuronal. We previously showed that sumatriptan administration over 6 days causes cutaneous allodynia, characterised by reduced periorbital and hindpaw withdrawal thresholds to von Frey, which subsides following withdrawal of the drug but can be re-evoked by exposure to a nitric oxide donor such as SNP. At day 20, sumatriptan, which has a half-life of approximately 2 hours (51), is no longer expected to be present in the circulation. The persistent susceptibility observed at this stage therefore most likely reflects a longer-lasting consequence of prior exposure rather than an acute pharmacological effect.

Chronic glial activation in the TNC at day 20, together with the delayed increase in Iba-1 and GFAP expression, may help explain how migraine-like responses can be more readily induced by known triggers following repeated sumatriptan exposure. The apparent change from predominantly neuronal pERK/pp38 association at day 6 to greater apparent glial association at day 20 is broadly consistent with this interpretation. However, because these colocalisation observations are based on representative images rather than formal quantitative analysis, they should be interpreted with caution. The present findings are therefore best interpreted as supporting a possible contribution of glial activation to maintenance of latent sensitisation rather than demonstrating a definitive shift from neuronal to glial drivers.

There is little evidence to suggest that 5-HT1B/1D receptors are expressed in glial cells of either the peripheral or central nervous system, whereas they are known to be expressed in trigeminovascular neurons. It is therefore unlikely that sumatriptan acts directly on glia to produce the activation observed here. A more plausible interpretation is that sumatriptan acts initially on neurons within the TGVS, which then indirectly promote glial activation through neuron-to-glia signalling. In response to a trigger such as an NO donor, these glial cells may in turn contribute to renewed neuronal activation and sensitisation. Thus, rather than demonstrating a definitive gliopathic mechanism of MOH, our data support the possibility that glial activation participates in the maintenance of latent sensitisation after repeated sumatriptan exposure.

### Sumatriptan-induced brain perfusion changes

Here, prolonged exposure to sumatriptan was associated with persistent changes in brain perfusion. Six days of sumatriptan exposure reduced cerebral blood flow compared with saline-treated rats, and this reduction remained evident at day 20 after withdrawal of the drug. In patients with migraine, changes in CBF during different phases of migraine attacks and during the interictal period have been extensively investigated using a range of approaches (29,52–53). These studies have raised a number of interpretive issues, particularly the inconsistency of CBF findings across studies. One explanation may be the inherently variable nature of migraine attacks, which differ in severity, duration and associated features, making comparison across patients and across separate attacks difficult.

In contrast, the present animal model shows relatively low variability, and by day 20 the animals are no longer exposed to sumatriptan and have returned towards normal sensory thresholds, thereby modelling a pain-free interval between attacks. In this context, the persistence of reduced CBF may represent a physiological correlate of the latent sensitised state. Although the MRI data in the present study are limited to CBF measures, they extend the behavioural and histological findings by suggesting that repeated sumatriptan exposure is associated with longer-lasting changes beyond the TG and TNC.

### Limitation to the study

Several limitations of the present study should be acknowledged. Subgroup sample sizes were modest, reflecting the complexity and intensity of the experimental design; however, statistically significant and internally consistent effects were observed across behavioural and molecular endpoints. Furthermore, power analysis was conducted to estimate the appropriate sample size. Additionally, all experiments were conducted in male rats. Given the higher prevalence of migraine in females and known sex differences in pain processing and neuroimmune signalling, future studies will be required to determine whether similar mechanisms operate in females and whether sex-specific differences influence susceptibility to medication-overuse headache. These considerations do not detract from the present findings but highlight important avenues for further investigation.

### Conclusions

Continuous sumatriptan administration in rats induces persistent alterations within the trigeminovascular system, including increased expression of biomarkers associated with neuronal activation and sensitisation in the trigeminal ganglion and trigeminal nucleus caudalis. Repeated sumatriptan exposure is also associated with delayed glial activation in the trigeminal nucleus caudalis, which may contribute to maintenance of latent sensitisation, although the underlying mechanisms remain to be defined. The persistence of reduced cerebral blood flow after drug withdrawal further indicates that repeated sumatriptan exposure is accompanied by longer-lasting changes in brain perfusion. Together, these findings support the value of this model for investigating the pathophysiology of medication-overuse headache and for informing translational studies of migraine chronification.

## SUPPLEMENTARY FIGURES

**S1. Representative dual-labelling of pp38 and Iba-1 in the trigeminal nucleus caudalis.** Images show saline- and sumatriptan-treated rats at day 6, day 20, and day 20 after SNP challenge. These panels qualitatively illustrate increased apparent association of pp38 signal with microglial profiles at later time points.

**S2. Representative dual-labelling of pERK and Iba-1 in the trigeminal nucleus caudalis.** Images show saline- and sumatriptan-treated rats at day 6, day 20, and day 20 after SNP challenge. These panels qualitatively illustrate increased apparent association of pERK signal with microglial profiles at later time points.

**S3. Representative dual-labelling of pp38 and GFAP in the trigeminal nucleus caudalis.** Images show saline- and sumatriptan-treated rats at day 6, day 20, and day 20 after SNP challenge. These panels qualitatively illustrate pp38 signal in regions of increased astrocytic labelling.

J.G.A.H. carried out the immunohistochemistry experiments, analysed the data. F.M.B. contributed to study design, supervised data analyses, and critically revised the manuscript. A.K. performed the MRI experiments, analysed imaging data, and contributed to data interpretation. M.D.F. conceived and supervised the study, obtained funding, contributed to experimental design and data interpretation, and revised the manuscript for important intellectual content. All authors approved the final version of the manuscript.

