## Supplementary figures and images for "Continuous sumatriptan exposure induces persistent trigeminovascular sensitisation and brain perfusion changes in a rat model of medication overuse headache"

### Supplemental figures

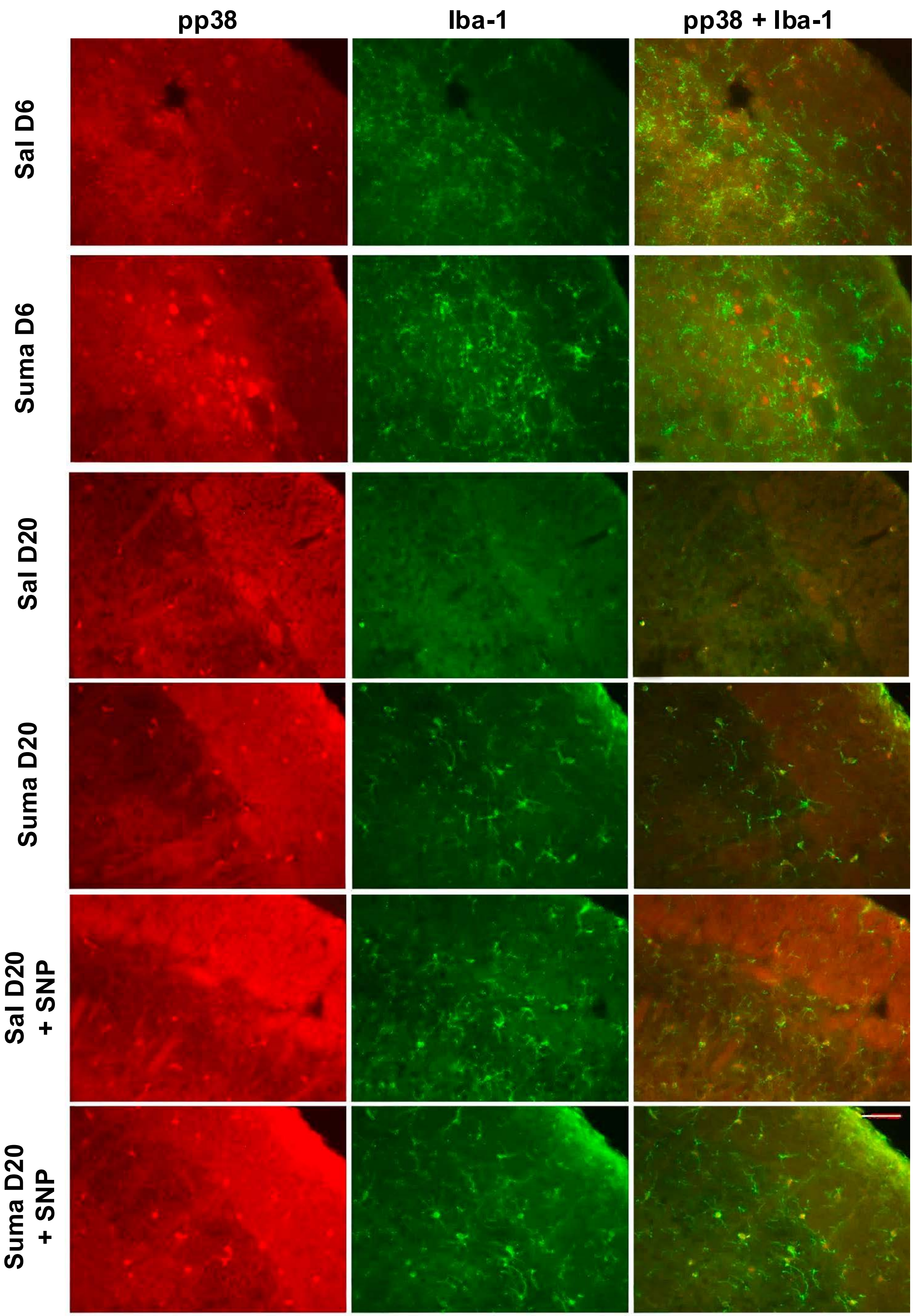

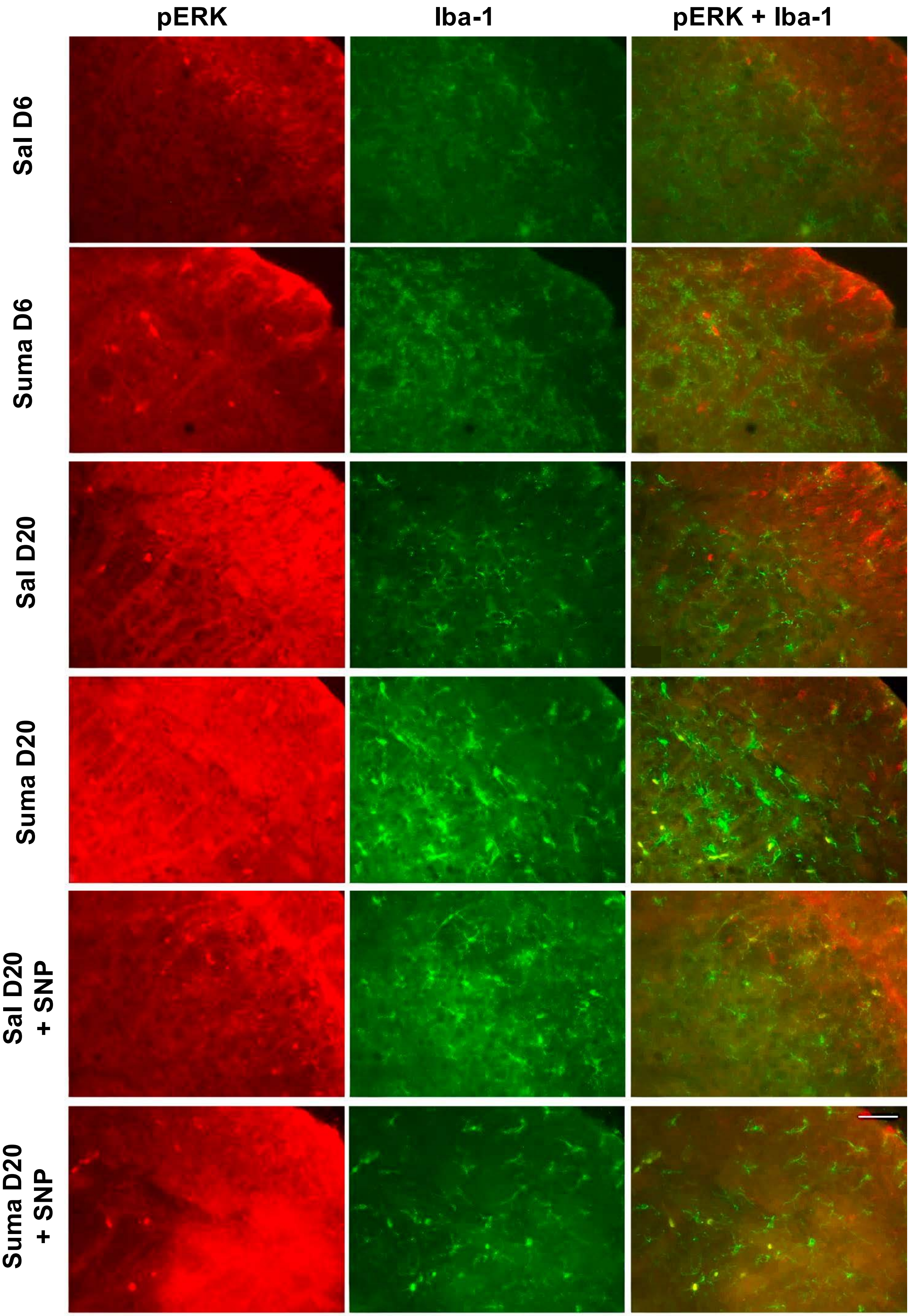

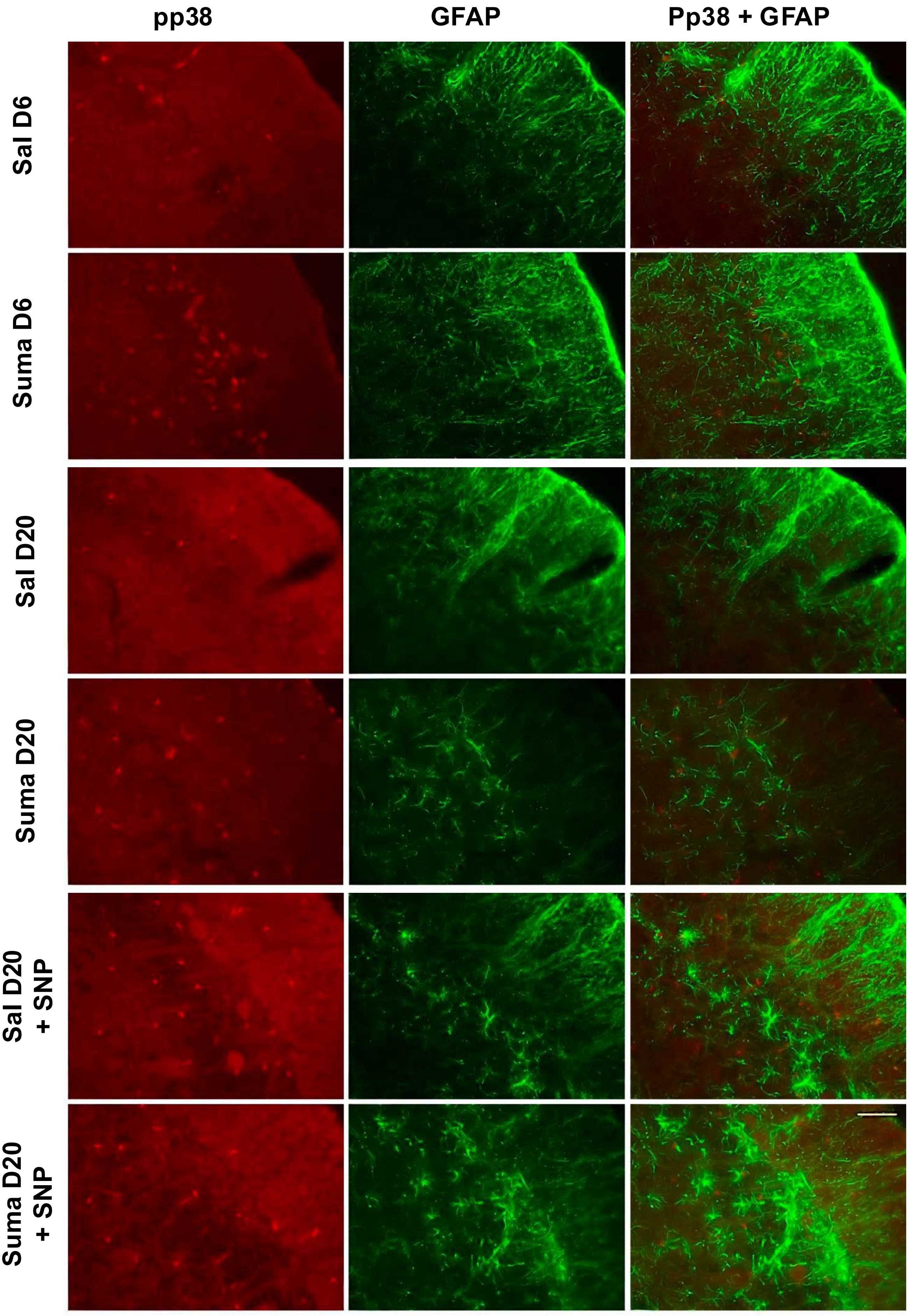
